# How citizen science could improve Species Distribution Models and their independent assessment

**DOI:** 10.1101/2020.06.02.129536

**Authors:** Matutini Florence, Baudry Jacques, Pain Guillaume, Sineau Morgane, Pithon Joséphine

## Abstract

Species distribution models (SDM) have been increasingly developed in recent years but their validity is questioned. Their assessment can be improved by the use of independent data but this can be difficult to obtain and prohibitive to collect. Standardized data from citizen science may be used to establish external evaluation datasets and to improve SDM validation and applicability. We used opportunistic presence-only data along with presence-absence data from a standardized citizen science program to establish and assess habitat suitability maps for 9 species of amphibian in western France. We assessed Generalized Additive and Random Forest Models’ performance by (1) cross-validation using 30% of the opportunistic dataset used to calibrate the model or (2) external validation using different independent data sets derived from citizen science monitoring. We tested the effects of applying different combinations of filters to the citizen data and of complementing it with additional standardized fieldwork. Cross-validation with an internal evaluation dataset resulted in higher AUC (Area Under the receiver operating Curve) than external evaluation causing overestimation of model accuracy and did not select the same models; models integrating sampling effort performed better with external validation. AUC, specificity and sensitivity of models calculated with different filtered external datasets differed for some species. However, for most species, complementary fieldwork was not necessary to obtain coherent results, as long as the citizen science data was strongly filtered. Since external validation methods using independent data are considered more robust, filtering data from citizen sciences may make a valuable contribution to the assessment of SDM. Limited complementary fieldwork with volunteer’s participation to complete ecological gradients may also possibly enhance citizen involvement and lead to better use of SDM in decision processes for nature conservation.

## Introduction

In the current context of biodiversity loss, a stronger relationship between conservation science and citizen participation could help to make conservation actions more effective (Ahmadi et al., 2017; Lewandowski & Oberhauser, 2017). Availability of data from citizen sciences has considerably increased over the past few decades (Dickinson, Zuckerberg & Bonter, 2010; McKinley et al., 2017). This data has great potential because (i) large quantities of data can be collected over large areas, which would be difficult and expensive for researchers to collect; (ii) data may be collected over long time periods, which is especially useful for studying the effects of climate and landscape changes on population dynamics at large scales; (iii) citizens are involved in the research process, thereby gaining knowledge, and their involvement might lead to improved implementation of biodiversity conservation action (Dickinson, Zuckerberg & Bonter, 2010; McKinley et al., 2017). However, quality of data from participatory sciences is heterogeneous and different methods have been developed to boost data accuracy and account for bias, including interactive project development, volunteer training, expert data validation and statistical modelling improvement (Kosmala et al., 2016). Although researchers have been skeptical about the value of datasets from citizen science, recent publications show that some could be as valid as data collected by professional scientists (Kosmala et al., 2016). This is conditional on such data being judged in context (i.e. according to the sampling methods used, program objectives and applications) on the use of rigorous data sorting and analyses (Isaac et al., 2014; Steen, Elphick & Tingley, 2019; Robinson et al., 2020).

Opportunistic presence-only data collected by citizens at large scales have contributed to the expansion of species distribution models (SDM) over the past twenty years, particularly for biological conservation applications (Guisan & Thuiller, 2005). The validity of presence-only SDM is however increasingly questioned as well as their applicability (Barve et al., 2011). Presence-only data come from different source databases reduced to simple species presence records and mostly collected in an unstandardized way by volunteers. In contrast to presence-absence data, they are abundant but have poor quality, few metadata and come from different sources (Robinson et al., 2020). This introduces numerous sources of bias that need to be assessed and accounted for in modelling processes (Phillips et al., 2009, Guillera-Arroita et al., 2015). Common problems are heterogeneous sampling effort, conditions and methods, imprecise spatial and temporal resolutions and different levels of expertise among observers (Schulman, Toivonen & Ruokolainen, 2007; Phillips et al., 2009, Dickinson, Zuckerberg & Bonter, 2010; McKinley et al., 2017). Different methods have been developed to correct these biases, including sorting or weighting presence-data to reduce identification errors and pseudo-replication linked to sampling effort (Guisan & Theurillat, 2000; Phillips et al., 2009) and/or using sampling effort assumptions to establish pseudo-absence sampling strategies (Barbet-Massin et al., 2012). Understanding the structure and intensity of sampling effort in space is essential to determine whether an undetected species is truly absent. For example, it may be conditioned by site accessibility (Kadmon, Farber & Danin, 2004; Phillips et al., 2009), site attractiveness or observer distribution (Phillips et al., 2009; Robinson, Ruiz-Gutierrez & Fink, 2018). Not accounting for heterogeneous sampling effort or using erroneous assumptions to define it can lead to over-assessment of model accuracy and/or false interpretation (Schulman, Toivonen & Ruokolainen, 2007; Phillips et al., 2009; Guillera-Arroita et al., 2015).

SDM validation is challenging (Vaughan & Ormerod, 2005) but is a crucial step for applying results to conservation objectives. There is still debate about SDM validity, especially when presence-only data is used to calibrate models (Barve et al., 2011). Using data with the same spatial bias to calibrate and assess a model tends to over-estimate prediction accuracy, by modelling observation processes more than ecological processes therefore producing erroneous results. Currently, testing model accuracy with a fully independent dataset is considered the most robust method for assessing SDM (Araujo et al., 2005; Guisan, Thuiller & Zimmermann, 2017). However, obtaining an external data set for large-scale studies is often cost prohibitive and exploiting data from standardized citizen science programmes may in some cases provide the solution. For example, Robinson et al. 2020 have shown that using filtered large-scale citizen science data for SDM calibration can improve model accuracy. Alternatively, detection-nondetection data from more standardized citizen sciences programs which are rarer than opportunistic data but have higher quality could provide presence-absence sets for external validation of presence-only SDM. In addition, using presence-only and presence-absence data at different stages of the modelling process could be a method for combining different data sets with heterogeneous quality which is a current challenge to improve SDM validity (Zipkin & Saunders, 2018; Robinson et al., 2020).

Amphibians are among the most threatened taxa in the world with rapid and widespread population declines (Stuart et al., 2004). They are particularly sensitive to fragmentation and habitat loss (Cushman, 2006) because they need different resources during their life cycle involving movements (seasonal migration and dispersion) between aquatic sites (usually ponds) and terrestrial areas (Sinsch, 1990; Cushman, 2006). Many citizen science programmes have been initiated for monitoring amphibian species (e.g. De Solla et al., 2005; Schmeller *et al*., 2008) and data collected have been used in some conservation studies to describe population trends (Petrovan & Schmidt, 2016), road effects (Cosentino et al., 2014), climate change (Préau et al., 2019) and large-scale species distributions (Brown et al., 2016). Despite abundant literature on amphibian ecology and the availability of several citizen science databases, few studies have attempted predictive amphibian distribution models at large scales (Graham & Hijmans, 2006; Brown et al., 2016; Préau et al., 2019). Therefore, amphibian data could be suitable for testing the capacity of different types of citizen data (presence-absence or opportunist) for calibrating and assessing SDM.

Here we compare the predictive performance of presence-only SDM for nine amphibian species using different types of data (*internal* presence-only or *external* presence-absence) from citizen science programs for their assessment. We also test the opportunity to use filtered standardized citizen science data to constitute the independent data set for external evaluation. We hypothesized that (1) the type of data used for validation (internal or external) would influence the assessment of model accuracy; (2) standardized citizen science datasets might be used as independent data for external evaluation of SDM using data filters and/or complementary fieldwork.

## Methods

### Study area

Our study was performed in Pays de la Loire (western France), a region covering 32 082 km^2^ with low relief and bordering on the Atlantic Ocean to the west. The region has an extensive hydrographic network organized around the River Loire and its tributaries, influencing local climate and landscape configuration. Agricultural landscapes dominate the region and traditional hedgerow network landscapes associated with extensive livestock farming are recognized for their conservation value. Such mosaics of small pastures delimited by hedgerows and small woods and generally associated with dense pond systems (Baudry, Bunce & Burel, 2000) are suitable for many organisms including endangered species such as some amphibians species (Boissinot, Besnard & Lourdais, 2019). With 21 known species (for 43 species recorded in France), the region has a high responsibility for the preservation of amphibians and their habitats, including traditional hedgerow landscape and wetlands.

### Biological data

We studied habitat suitability of 9 amphibian species: *Bufo spinosus, Hyla arborea, Rana dalmatina, Rana temporaria, Triturus cristatus, Triturus marmoratus, Lissotriton helveticus, Salamandra Salamandra*, and *Pelodytes punctatus*. Two types of amphibian data were used: (1) opportunistic data from a citizen database with presence-only records for model calibration and internal evaluation (2) standardized detection-nondetection data from a citizen science programme and complementary field work for external evaluations. A more detailed description of the data sets and complementation strategies is available in Appendix 1.

#### Opportunistic presence data (calibration and cross-validation dataset)

We accessed presence-only occurrences from a regional database for the period 2013-2019. 86% of the dataset was collected by citizens and recorded online (website or associated mobile application) and 14% by various professional organisations involved in nature protection. All data were compiled for the regional Atlas of amphibians by a French non-governmental organisation (French BirdLife partner - LPO). See Appendix 1 Table 1 for data sources. We retained only species with enough data according to number of predictors used (i.e. at least 477 presence cells; see Appendix 1 Table 2). We selected only precise GPS records (precision of the observation under 50 meters) and we checked all data for anomalies in geographical location or species identification.

For each species, we sorted data to reduce spatial autocorrelation by projecting presence data on a 500m resolution grid and retaining only cells containing at least one occurrence as presence cells for the analyses (Guisan & Theurillat, 2000). We chose a 500m resolution as it is the mean size of species’ home ranges (Semlitsch & Bodie, 2003). Finally, we excluded all opportunist data intersecting cells used for external validation described below to increase independency between calibration and validation sets.

#### Standardised detection-nondetection data (external validation datasets)

For external validation, we firstly extracted *detection-nondetection* amphibian data for the period 2013-2019 collected as part of a citizen science program called “Un Dragon dans mon Jardin” (Appendix 1 section 1.2.). We retained 576 sites which were monitored at least 3 times between February and June during at least one year and following a standard method commonly used for amphibian community surveys (Boissinot, Besnard & Lourdais, 2019). We called this dataset *CS.0* (see Figure 1). Some large areas of the region were not sampled due to lack of observers so that data were clustered near cities, with spatial autocorrelation. Therefore, with help from volunteers, we completed this dataset with some additional fieldwork and applied filters.

**Figure 1.**
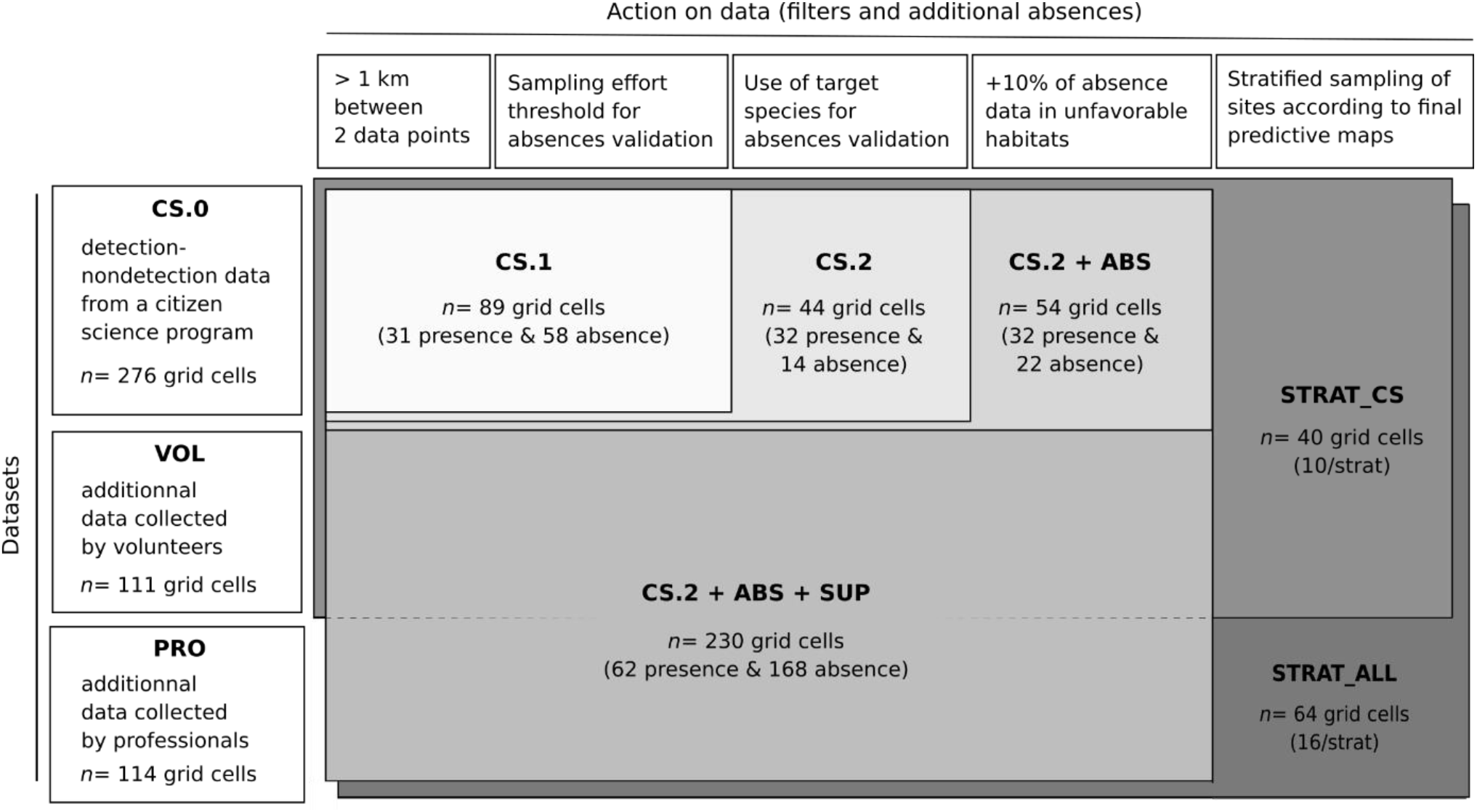
Description of datasets, filters and complementation used for external evaluation derived. Number of data (*n* grid cells) given for *T. marmoratus*, for 1 iteration only and s3. CS: data derived from the citizen science data set; SUP: additional data collected by volunteers (VOL) and professionals (PRO).

To complete and filter *CS.0*, different strategies were used. Firstly, we organised complementary fieldwork in 2018 and 2019 to complete two landscape gradients (woody element density and pond density) which are two variables known to strongly affect amphibian distribution and which are relevant in our regional context. All sites were selected randomly but so as to maximise and decorrelate the two landscape gradients in different areas (see Appendix 1 section 1.3.). In total, 263 sites were monitored: 132 sites by experts in 2018-2019 (called *PRO*, see Figure 1) and 131 by 75 volunteers in 2019 (called *VOL*, see Figure 1). All data (*CS.0*, *PRO* and *VOL*) were projected on the same 500m resolution grid. One further problem, common in citizen science programmes (Geldmann et al., 2016), is that only aquatic sites are surveyed while areas known to be very unsuitable for amphibians such as urban areas and intensive agriculture are generally excluded. To reduce this source of bias, we randomly selected 5% more 500m grid cells in totally urbanized areas without aquatic sites and 5% more grid cells in homogeneous croplands without trees or ponds and we attributed “absence” values to each after field checks (called *ABS*, see Figure 1). These landscapes represent 9% of the total area of the region.

Secondly, we applied different filter combinations to establish sub-sets from *CS.0, PRO* and *VOL* (Figure 1):

1. a minimum distance of 1km between two grids containing data;
2. threshold values to validate non-detection as absence data and exclude under-sampled sites, defined as a minimum sampling effort required to detect a species based on the species’ detectability group and observer level of expertise. Four species detectability groups were defined from occupancy studies in France (Boissinot, 2009; Petitot et al., 2014) and the UK (Sewell, Beebee & Griffiths, 2010). Observers were classed as either *expert, intermediate* or *novice* using 3 criteria: number of years of participation, number of species observed and permit-holder for amphibian capture. A *“novice”* observer was considered more likely to miss or misidentify a species which was actually present than an *“expert”* observer for the same considered survey effort. In addition, novice observers did not use sampling nets, influencing detectability, in particular of Urodeles. Based on our observer classes and sampling methods used (e.g. acoustic, visual and/or fishing), we set threshold values for the minimal sampling effort needed to validate absence data, depending on species detectability (see Appendix 1 section 1.4. for details) and according to the results from Boissinot 2009 for minimum sampling effort needed to detect a focal species (with 95% probability) in a similar biological and landscape context;
3. target species to valid nondetection as absence, as recommended by Phillips and co-workers in 2009. So, if *species A* with the same detectability as *species B* is detected at a site, then *species B* is likely to be truly absent (see Appendix 1 section 1.5. for target species list);
4. stratified sampling on final prediction maps (see STRAT_CS and STRAT_ALL in Figure 1). We established independent datasets, stratified by model predictions for each species (Newbold et al., 2010; Guisan, Thuiller & Zimmermann, 2017) with equal random sampling of validation cells with presence or absence data in 4 habitat suitability classes predicted (i.e. [0;0.3[, [0.3;0.5[, [0.5;0.7[and [0.7;1]). We obtained an equal number of validation cells by predicted suitability class (see 1.4.3. for predictive map used).

### Environmental dataset and variables

We assembled environmental data relevant to amphibian ecology and of importance in the study region (Guisan, Thuiller & Zimmermann, 2017). A more detailed description of variables with associate references is available in Appendix 2 Table 1.

Bioclimatic variables were accessed from a compilation of climate data for the period 1950-2000 at a spatial resolution of 5km2 (Hijmans et al., 2005). An altitude variable was derived from the U.S. Geological Survey’s Hydro-K data set, at the same spatial resolution. We performed a principal-components analysis (PCA) on 11 bioclimatic variables relevant for amphibians and the altitudinal layer to produce 2 uncorrelated axes (see Appendix 2 Table 4 and Figure 2). Land cover data were downloaded from the highly detailed vector database OCS GE 1.1 (IGN 2019), the Theia OSO Land Cover Map 2017 (available at www.theia-land.fr) and from BDTopo (IGN 2019). This was coupled with a more detailed regional inventory of hedgerows (from 2005 to 2008) and ponds (2012) and a national farming database from the EU LPIS (Land Parcel Identification System 2016) used to classify agricultural areas (see Table 1).

**Table 1.** Environmental variables used for species distribution modelling of each amphibian species in Pays de la Loire region. Associated references are available in the Appendix 2 Table 1.

| Variable category | Code | Variable description | Original resolution |
| --- | --- | --- | --- |
| Climatic | CLIM_1 | first axis from a PCA on 12 worldclim variables and altitude | 2.5 arc-min/5km |
|  | CLIM_2 | second axis from a PCA on 12 worldclim variables and altitude | 2.5 arc-min/5km |
| Land cover | %WOOD_DM | proportion of deciduous and mixed forest | 5m |
|  | %WOOD_C | proportion of coniferous forest | 5m |
|  | %CROP | proportion of crop | 20m |
|  | %PASTURE | proportion of permanent pasture | 20m |
|  | NB_PONDS | pond density (or water point density) | 5m |
|  | L_HEDGE | hedgerow density | 5m |
|  | L_ROAD_1ST | primary road density outside urban areas | 5m |
|  | L_ROAD_2ND | secondary road density outside urban areas | 5m |
|  | L_RIVER | canal and river density | 5m |
|  | %URBAN | proportion of urban area | 20m |

We calculated land-cover variables in windows composed of a 500m grid cell with a buffer of 300m (see Table 1). This took into account landscape context based on species’ dispersal capacities as well as the resolution of the species data set (Guisan & Thuiller, 2005). Distance and home range differ among amphibian species but a 1km circle may be accepted as an average maximum range (Collins & Fahrig, 2017). Collins and Fahrig (2017) and Boissinot et al. (2019) show that landscape variables affect Anuran occupancy and diversity at this scale in agriculture-dominated regions. We use the same environmental variables for all species (see Table 1) except *B. spinosus* for which “pond density” (water point <5000m2) was substituted by “water point density” because of this species’ ability to reproduce in larger water bodies with fish (Boissinot, Besnard & Lourdais, 2019).

All predictive variables were centred and scaled. The spatial correlation between environmental predictors was investigated using the variance inflation factor (VIF) as a measure of multicollinearity and Pearson correlation tests with VIF<6 and r<0.6 as advised by O’Brien, 2007 (see Appendix 2 Table 2 and 3).

### Habitat suitability modelling

#### Statistical models

Different modelling algorithms can lead to varying results according to heterogeneous sensitivities and calculation processes (Thuiller et al., 2009). Therefore, consensus models based on multi-modelling approaches (ensemble-modelling) can improve final results by reducing ‘noise’ associated with individual model errors (Araujo et al., 2005; Thuiller et al., 2009; Meller et al., 2014). For each species, we used one regression-based approach (Generalized Additive Models, GAM) and one machine learning algorithm (Random Forest, RF) to predict and assess habitat suitability within the studied region with 50 bootstrap replicates. Presence points were randomly split 50 times into a training set (70% of the whole dataset) and the remaining 30% were used as testing set for internal evaluation (see 1.4.). To construct these models, we used biomod2 package (Thuiller et al., 2009) in R environment v. 3.5.3 (R Development Core Team, 2019).

#### Background data and pseudo-absence selection

Modelling habitat suitability for a species with GAM or Random Forest requires both presence and absence data. In order to overcome the problem of missing absence data needed for most SDM, pseudo-absence selection strategies have been developed to select absence data where real absence is most likely (Barbet-Massin et al., 2012, Phillips et al., 2009). We tested three strategies for generating artificial absence points: (s1) simple random selection of background points within the studied region (Guisan, Thuiller & Zimmermann, 2017); (s2) random pseudo-absence selection excluding known presence points (Engler, Guisan & Rechsteiner, 2004); (s3) random pseudo-absence selection constrained to take sampling effort into account (see Appendix 3 for method). The latter strategy aimed to select pseudo-absences where true absences were more likely. For this strategy we considered three main sources of bias in pseudo-absence selection: accessibility, linked to distance from roads or urban areas (Kadmon, Farber & Danin, 2004; Barbet-Massin et al., 2012), attractiveness, relating to oversampling in protected sites or tourist areas (Phillips et al., 2009; Robinson, Ruiz-Gutierrez & Fink, 2018) and observer effort, because certain administrative areas are covered by particularly active nature protection organisations (see Appendix 3). For each strategy, the number of artificial absences was fixed equal to the number of presence data (Barbet-Massin et al., 2012; Liu, Newell & White, 2019) and we performed 10 replicates of the artificial absence generation processes.

#### Ensemble modelling

Finally, we conducted ensemble modelling by calculating the median value of (1) all individual maps generated by GAM and Random forest (i.e. 500 maps/algorithm) (Thuiller et al., 2009) to compare internal *versus* external evaluation for each species. Secondly, we also calculated median values from ensemble maps calibrated with 100% of presence-only data to compare different external evaluations sets (i.e. 10 maps/algorithm).

### Internal and external model validation

We firstly use a cross-validation method using a 30% random split of the whole set to asses each model (for pseudo-absence selection strategies s1, s2 and s3) with 50 bootstraps repeated 10 times. We calculated the area under the curve (AUC) of a receiver operating characteristic (ROC) plot of the predicted model habitat suitability scores with (1) the 30% test dataset for internal validation and (2) with the larger filtered external independent dataset (e.g. *CS.2+ABS+SUP*, see Figure 1) using Biomod2 package (Thuiller et al., 2009). AUC is the most common metric used in SDM studies, as it has the advantage of being threshold and prevalence independent and has been accepted as the standard measure for assessing SDM accuracy (Guisan, Thuiller & Zimmermann, 2017). AUC > 0.50 signifies that the model has better prediction than a random model. Secondly, we calculated AUC values, specificity (true negative rate) and sensitivity (true positive rate) of ensemble models calibrated with 100% of the presence-only data using different evaluation sets derived from the global external dataset used in the previous stage (see Figure 1). These calculations (with 100 bootstraps) were performed using PresenceAbsence package (Freeman 2012) with a standard threshold value for presence-absence discrimination fixed at 0.5.

## Results

### Model performance and selection

For each species, the median AUC was higher with internal validation than external validation for all three pseudo-absence selection strategies (s1, s2 and s3), both for GAM and Random Forest (see Figure 2 and Appendix 4) with a delta-AUC ranging from 0.05 (*T. marmoratus*) to 0.21 (*B. spinosus*). With internal evaluation (cross-validation), all models had excellent (AUC>0.90) very good (0.80-0.90) or good accuracy (0.70-0.80) except for the model of *B. spinosus* and *H. arborea* including sampling effort parameters (s3). However, with external evaluation, only four species had a high level of accuracy (AUC>0.70): *S. salamandra, T. marmoratus, P. punctatus* and *R. temporaria*. For *R. dalmatina, T. cristatus* and *L. helveticus*, model accuracies were poor (0.60<AUC<0.70) and for *B. spinosus* and *H. arborea* even poorer (AUC<0.60). The strategy s1 (background data) was not selected neither with internal nor external evaluation.

**Figure 2.**
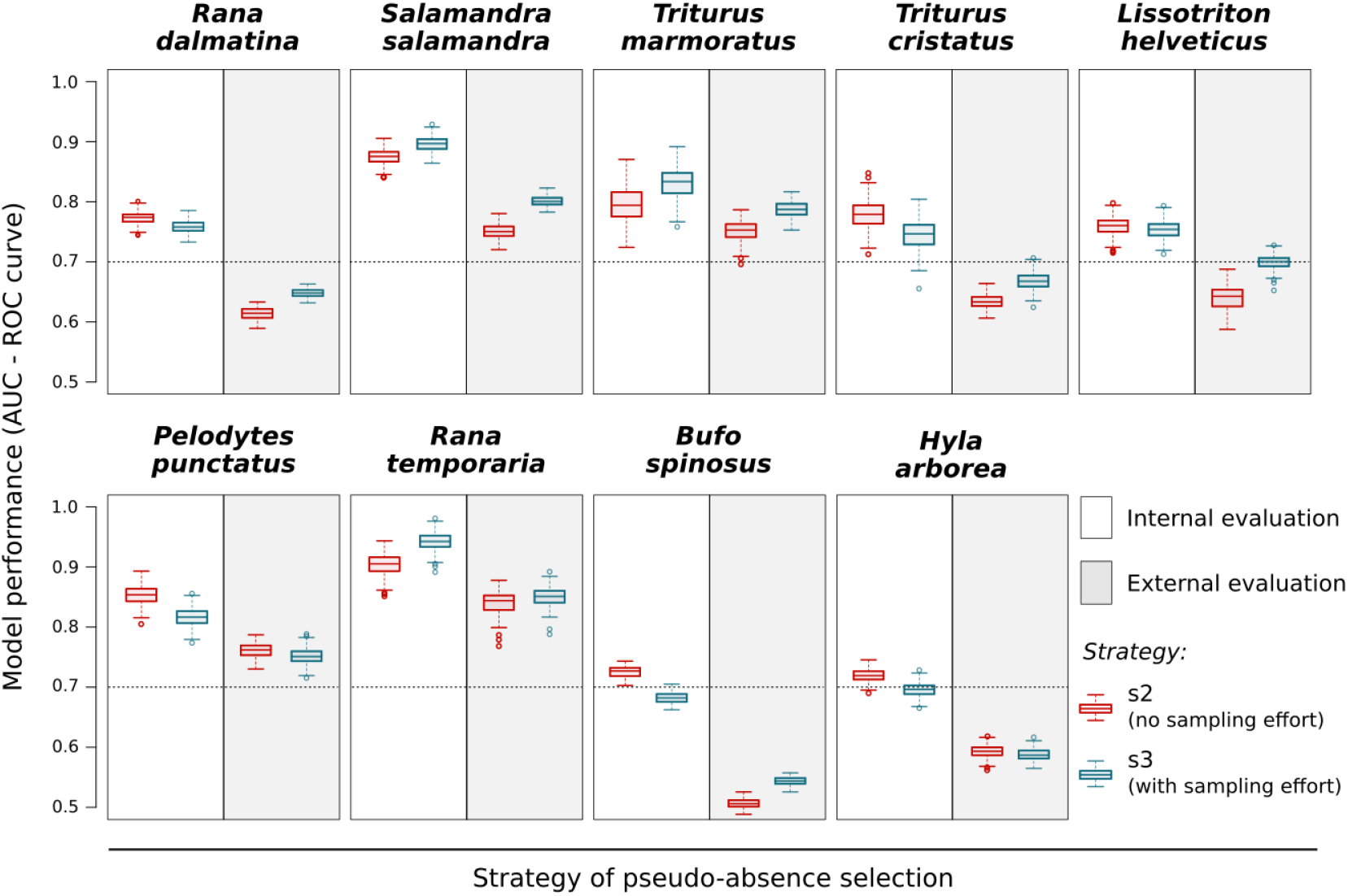
Model performance for the 9 studied species assessed by external or internal data using different pseudo-absence selection strategies. Assessment by AUC under the ROC for GAM only are shown (see supplementary material). Artificial absence sampling strategies shown are s2 (random pseudo-absence selection excluding known presence points) and s3 (random pseudo-absence selection excluding known presence points and adjusted to consider site accessibility and sampling effort). Per strategy, 10 replicates of the artificial absence points generation processes with 50 bootstraps for the random selection of the straining set (70%) and the internal testing set (30%). Black dotted line indicates the 0.70 threshold above which models have an acceptable level of accuracy (Swets 1988).

**Figure 3.**
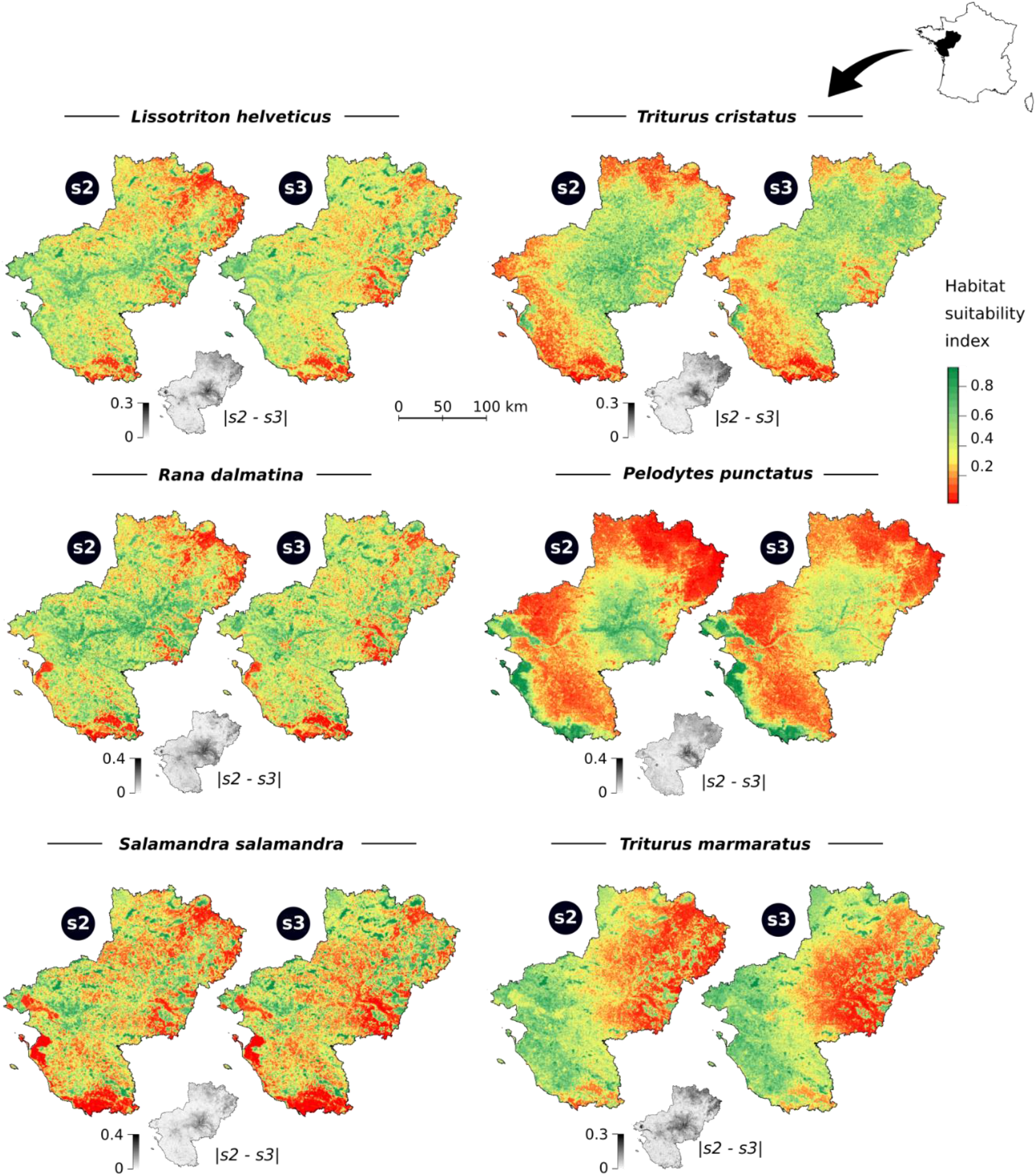
Habitat suitability maps for six studied species produced using two form of pseudo-absence selection. s2 (random pseudo-absence selection excluding known presence points) and s3 (random pseudo-absence selection excluding known presence points and constrained to account sampling effort). The black and white map under each pair shows net difference between s2 and s3. Map resolution is 500m.

The method used for pseudo-absence selection influenced the predictive performance of models but differences between AUC values were minimal (Figure 2). However, s2 (uncorrected sampling bias) was the best strategy for six species when internal validation was used, while s3 was best for seven species when external validation was used. Results for RF can be found in the supplementary material but do not differ greatly (Appendix 4 Figure 1).

### Impact of model selection on final habitat suitability map

Internal or external validation resulted in different models being selected, based on AUC comparison. Therefore, the final habitat suitability maps selected by each of these two assessment methods would lead to different interpretation and conservation decisions. Maps for *H. arborea* and *B. spinosus* are not shown because of poor accuracy (see supplementary results Appendix 4). All response curves and associated variable contributions can be found in the supplementary material (Appendix 4 Figures 2 and 3).

### Comparison of external evaluation sets

Values of AUC, sensitivity and specificity to four species are shown in Table 2 (two Anurans and two Urodeles; one forest species and one generalist specie each). Results for other species and CS.1 (similar to CS.2) are presented in Appendix 4 Table 3. Considering AUC values, evaluation with the external dataset from participative science without filter data (*CS.0*) show more similar model selection results than internal cross-validation except for *R. temporaria*. Sorting presence data led to decreased sensitivity and increased specificity for all species except for *S. Salamandra*. The models selected (s2 or s3) were similar for most species whether using stratified data from volunteers’ only (STRAT_CS) or stratified data with added professional observations (STRAT_ALL), or professional data only (PRO). See Table 2 and Appendix 4. We excluded the s1 model from the comparison because this model is never selected, either with internal or external evaluation.

**Table 2.** Model performance according to different filters and complementary fieldwork applied to the external evaluation dataset. External datasets used were (see Figure 1): *CS.0* (all data from the standardized citizen science dataset); *CS.2 + ABS (CS.2* with 10% supplementary absence cells in very unsuitable habitats); *PRO* (data collected by professionals only in 2018-2019); *CS.2 + ABS + SUP* (citizen science data cited before adding all complementary fieldwork by professionals and volunteers); STRAT_CS (stratified data selection from CS.2+ABS with complementary fieldwork by volunteers); STRAT_ALL (stratified data selection from CS.2+ABS+SUP). Models assessed: s2 (random pseudo-absence selection excluding known presence points) and s3 (random pseudo-absence selection constrained to account sampling effort and correct sampling bias). SEN: sensitivity, SPE: specificity. Bold values show best values between s2 et s3 with delta > 0.02 and grey cells show species with less than 2 presence data. All analyses with a random sampling in presence selection with a distance condition or a stratified random selection were performed using 100 bootstraps (mean calculation).

|  |  | *Triturus marmoratus* |  |  | *Lissotriton helveticus* |  |  | *Hyla arborea* |  |  | *Rana temporaria* |  |  |
| --- | --- | --- | --- | --- | --- | --- | --- | --- | --- | --- | --- | --- | --- |
|  |  | SEN | SPE | AUC | SEN | SPE | AUC | SEN | SPE | AUC | SEN | SPE | AUC |
| CS.0 | s2 | 0,63 | 0,41 | 0,58 | **0,69** | 0,29 | 0,53 | **0,82** | 0,43 | 0,68 | 0,71 | **0,87** | 0,87 |
|  | s3 | **0,65** | **0,45** | **0,61** | 0,64 | **0,34** | 0,54 | 0,78 | **0,49** | 0,68 | **0,86** | 0,84 | 0,88 |
| CS.2 + ABS | s2 | 0,59 | 0,78 | 0,81 | **0,67** | 0,76 | 0,80 | **0,71** | 0,61 | 0,73 | 0,64 | **0,88** | 0,86 |
|  | s3 | 0,58 | **0,87** | 0,81 | 0,62 | **0,88** | **0,86** | 0,68 | **0,64** | 0,74 | **0,82** | 0,84 | 0,86 |
| CS.2 + ABS + SUP | s2 | 0,58 | **0,82** | 0,82 | **0,51** | 0,77 | 0,68 | **0,63** | **0,56** | 0,63 | 0,59 | **0,91** | 0,85 |
|  | s3 | **0,62** | 0,78 | 0,81 | 0,49 | 0,78 | **0,74** | 0,57 | 0,54 | 0,63 | **0,76** | 0,86 | 0,85 |
| PRO | s2 | 0,58 | **0,83** | **0,85** | **0,41** | 0,76 | 0,61 | **0,59** | **0,39** | 0,51 | 0,50 | 0,95 | 0,71 |
|  | s3 | **0,79** | 0,77 | 0,83 | 0,37 | **0,79** | **0,64** | 0,51 | 0,33 | **0,54** | 0,50 | 0,92 | 0,71 |
| STRAT_CS | s2 | 0,71 | **0,72** | **0,81** | 0,67 | 0,81 | 0,82 | 0,72 | 0,64 | 0,73 | 0,79 | 0,57 | 0,69 |
|  | s3 | 0,70 | 0,69 | 0,78 | 0,68 | 0,82 | **0,88** | 0,71 | 0,63 | **0,75** | **0,95** | **0,60** | **0,79** |
| STRAT_ALL | s2 | **0,80** | **0,74** | **0,87** | 0,66 | 0,76 | 0,78 | **0,68** | **0,62** | 0,70 | 0,84 | 0,57 | 0,73 |
|  | s3 | 0,78 | 0,70 | 0,83 | **0,71** | **0,83** | **0,89** | 0,66 | 0,60 | **0,72** | **0,96** | 0,58 | **0,81** |

## Discussion

External evaluation with independent data generated lower AUC values than cross-validation, which calls into question the validity of models validated by commonly used selection threshold values such as AUC> 0.70. According to Araujo et al. 2005, internal evaluation with non-independent data always leads to over-optimistic assessment of model performance. Even if cross-validation is better than substitution procedures (Araujo et al., 2005; Vaughan & Ormerod, 2005; Edwards et al., 2006), split data for internal validation are non-independent and do not avoid the main limits of correlative models in SDMs because of spatial or temporal autocorrelation, especially when sampling effort is highly heterogeneous (Edwards et al., 2006, Roberts et al. 2017). Our result supports criticisms of certain types of SDM and further highlights the need to be careful in their general interpretation and assessment (Lobo, Jiménez-valverde & Real, 2008).

The difference between internal-AUC and external-AUC is particularly pronounced for the most common and generalist species in spite of the large number of available data, especially for *B. spinosus* and *H. arborea* whose models failed to attain an acceptable level of accuracy with external evaluation. Brotons et al. 2004 highlighted the difficulty of predicting distributions of the most generalist species. However, for such species, the use of filters increases specificity values considerably and the results are coherent with these species’ wide ranging and ubiquitous distributions. Using external presence-absence data also makes it possible to exploit the whole presence-only data set for calibration and to use stricter filters to reduce sampling bias or data culling to retain higher quality data (Steen et al. 2019, Isaak et al. 2014). Our study shows that it is possible to apply strong filters (e.g. STRAT_CS) but finally retain reasonable sample sizes for most species.

It should be noted that, for four species, *R. temporaria, T. marmoratus, S. salamandra* and *P. punctatus*, our results were ambiguous. For the first three, all forest-dwelling species or very closely related to woodlands (Boissinot, Besnard & Lourdais, 2019), both internal and external validation methods selected models with sampling effort integrated (s3). Two main reasons could explain these results: firstly, presence data may have been insufficient for *R. temporaria*, and secondly these species’ affinity for forest habitats. *R. temporaria* is a rare species and is more dependent on wet forest, flood meadows and small streams as breeding sites than other species (Boissinot et al., 2015). This species has a patchy distribution (i.e. locally abundant but regionally rare) and is difficult to detect. Hence, presence data are few in the studied area both in the opportunist dataset (N=477 presence-cells) and in the independent dataset (N=13 presence-cells). As highlighted by Vaughan & Ormerod (2005) such factors can lead to model over fit, even with a relatively small number of variables, resulting in high AUC values. In addition, according to Brotons et al. (2004), low-density habitat (i.e. forest habitat in our region) may be over-weighted and it can be difficult to assess between good or bad suitability without adapted presence-absence data. Monitored forest sites are rare in our validation dataset and the assumptions we used to define sampling effort may not be well adapted for forest specialists. Finally, *P. punctatus* is a rare species but abundant on the Atlantic coast. Unlike the other species, it is a pioneer, adapted to open areas, especially primary unvegetated habitats such as sand dunes and mudflats with frequent physical disturbance (Joly et al., 2005). These habitats are mainly located near the coast and along the main regional floodplains (Loire Valley), with a high local density of presence data. So this species is also patchily distributed and models may be affected by the same bias as *R. temporaria*. Alternatively, the similarity between AUC values may also relate to sampling effort bias along the Atlantic coast (e.g. Fithian et al., 2015). There results highlight the need to adapt methods and filters used for each species.

### Using heterogeneous data from citizen science in SDM

Our results show that it is possible to obtain useful external and independent datasets for model validation from filtered standardized citizen science data. Indeed, the use of filters have successfully reduce bias and noise in citizen science data sets for SDM in others studies (Robinson, Ruiz-Gutierrez & Fink, 2018; Isaac et al., 2019; Steen, Elphick & Tingley, 2019). In addition, filtered evaluation dataset showed coherent results according to Phillips et al. 2009. Indeed, choosing pseudo-absence data with the same bias as occurrence data improved model performance.

Since external independent data is necessary for more robust assessment of SDM (Araujo et al., 2005), but prohibitive to collect, filtering low quality but large datasets from monitoring to obtain more standard and independent data may be worthwhile. In addition, AUC appears to be more informative when presence-absence data is used to assess and compare models than when presence-background data alone is used (Jiménez-Valverde, 2012). However, using detection-nondetection citizen data without filters may also lead to erroneous results because of overlapping sources of bias in both datasets (e.g. *CS.0* selects the same model as cross-validation). The large amount of available data allows us to strongly select data according to our research objective. Our results using *PRO* datasets are inconclusive for rare species perhaps because their detection was insufficiently frequent (e.g. only two observations of *R. temporaria* for 132 sampled sites). Finally, we found that general rules to guide data sorting were difficult to define. Our results were sensitive to the type of data used, and the species studied, reinforcing the need to define filters according to available data and species’ ecology (Steen, Elphick & Tingley, 2019).

Independence between training and testing sets is an essential criterion, but data should also be unbiased or corrected. Selection methods have been developed to try to divide the opportunistic dataset strategically to increase the independence of the testing set for cross-validation (see Block-cross-validation in Robert et al. 2017). However, this method does not make it possible to escape from the general biases linked to sampling effort and/or can create extrapolation problems (see Robert et al. 2017). Using a more standardized dataset from a participatory science program (e.g. CS0) for the evaluation makes it easier to understand the sources of bias (presence of metadata and non-detection data), to better control them and to obtain more robust information on the absence data. However, these data may also share biases with the opportunistic dataset used for calibration. In our case, the sampling of the monitored sites (*CS0*) was partly biased towards volunteers’ place of residence and areas with a higher density of observers. These biases were reduced through additional sampling involving volunteers. Our results show that certain filters, as well as targeted complementary fieldwork, make it possible to reduce the biases identified and produce conclusive results. In addition, the use of a stratified sampling of the testing set along the suitability gradient from the SDM results (e.g. our STRAT_CS and STRAT_ALL datasets) appears to be a particularly interesting method showing stable and consistent results according to Phillips et al 2009.

However, our method may be not applicable in all cases. In our study, external data came from a program with general population monitoring objectives, using standard methods designed to be accessible to a wide audience (eg. novice and professionals). This program concerns all amphibians and their habitats, whereas many citizen science programs are limited to a particular taxonomic group or habitat type (cities, gardens or farms…) and would therefore be difficult to extrapolate to wider contexts.

### Involved stakeholders and citizens in conservation research

Our study was part of a wider project for amphibian conservation in the Pays-de-la-Loire region of western France. Involving citizens in the SDM evaluation process may make conservation action easier to implement, through both better shared knowledge and stronger personal involvement. Forrester et al. (2017) and Lewandowski and Oberhauser (2017) highlighted an increase in conservation advocacy among participants of citizen science projects that might improve access to evidence for conservationists and decision makers (Sutherland & Wordley, 2017). Maps are a specially a good tool for improving communication between researchers and volunteers in the context of citizen science (Zapponi et al., 2017). Indeed, many nature protection organisations are already involved in distribution atlas projects and naturalists are aware of data collection methods and local species distributions. They seek out ways to prioritise field observations; making a useful contribution to developing SDMs to guide conservation action can be a source of motivation, making scientist-volunteer interactions easier.

## Data accessibility

Data sample and access procedure are available online: https://doi.org/10.5281/zenodo.4043460

## Supplementary material

Script and codes are available online: https://doi.org/10.5281/zenodo.4043460

## Acknowledgements

This work would not have been possible without the support of the Pays de la Loire Herpetological Group, the CPIE Regional Union and the French BirdLife International partner (LPO). We are especially grateful to Benoit Marchadour (LPO) who coordinates the regional distribution atlas. We also acknowledge the many naturalists involved, for access to data and for additional fieldwork, especially Dorian Angot, Anne-Lise Charpentier, Baptiste Gaboriau, Ludovic Aubry, Jeanne Abbou, Flore Gamet, François Varenne, Adrien Maitrepierre, Leslie Michal, and Martin Bonhomme. We thank Xavier Lozupone for his help for mapping of environmental variables at regional level and Véronique Beaujouan, Adeline Bulot, Hervé Daniel, Gilles Martel and Aurélien Besnard for providing helpful discussion. Our work was supported by funding from Angers Loire Metropole, The French Society for Ecology and Evolution and “Humanity and Biodiversity”.

Version 4 of this preprint has been peer-reviewed and recommended by the *Peer Community In Ecology* (https://doi.org/10.24072/pci.ecology.100059)

## Appendix

Appendix 1-4: https://www.biorxiv.org/content/10.1101/2020.06.02.129536v4.supplementary-material

