## Appendix 1 for "How citizen science could improve Species Distribution Models and their independent assessment"

### Appendix 1 – Data description and complementation

#### 1.1. *Opportunistic presence data (calibration and cross-validation dataset)*

| Database name | Type | General website | Data proportion |
|---|---|---|---|
| Faune_anjou | Citizen bases with validation process by professionals | https://www.faune-anjou.org/ | 25% |
| Faune_maine | | https://www.faune-maine.org/ | 10% |
| Faune_vendee | | https://www.faune-vendee.org/ | 10% |
| Faune_loire_atlantique | | https://www.faune-loire-atlantique.org/ | 11% |
| Biolovision | | https://data.biolovision.net/ | 18% |
| URCPIE | Professional & volunteers | http://urcpie-paysdelaloire.org/ | 12% |
| Bretagne Vivante | Naturalist group | https://www.bretagne-vivante.org/ | 2% |
| ONF_BDN | Professional | https://www.onf.fr/ | 3% |
| SICEN | Professional | http://www.cenpaysdelaloire.fr/ | 3% |
| BASEPARC PNRMP / OPN | Professional | https://pnr.parc-marais-poitevin.fr/ | 2% |
| Naturalistes en lutte | Naturalist group | https://naturalistesenlutte.wordpress.com/ | 2% |
| Sterne 2.0 | Professional | http://www.sterne2.com/ | 1% |
| Les naturalistes vendeens | Naturalist group | http://naturalistes-vendeens.org/ | 1% |
| Gouret_FLA | Naturalist individual base | - | <1% |
| Cap Atlantique | Professional | https://www.cap-atlantique.fr/accueil | <1% |
| Undragon.org | Citizen base | http://undragon.org/ | <1% |
| ONCFS | Professional | http://www.oncfs.gouv.fr/ | <1% |

**Table 1. Data sources**

General coordination of the regional Atlas of amphibians: Ligue pour la Protection des Oiseaux – Pays-de-la-Loire (http://paysdelaloire.lpo.fr/).

| | Opportunistic presence-only dataset (model calibration and cross-validation) | |
|---|---|---|
| **Species** | Total nb of presence | Nb of 500m presence-cells |
| **Anourans:** | | |
| *Bufo spinosus* | 8320 | 4127 |
| *Hyla arborea arborea* | 6344 | 3353 |
| *Pelodytes punctatus* | 2711 | 1103 |
| *Rana dalmatina* | 9073 | 3752 |
| *Rana temporaria* | 1525 | 477 |
| **Urodeles:** | | |
| *Salamandra Salamandra terrestris* | 4916 | 2242 |
| *Triturus marmoratus* | 1478 | 629 |
| *Triturus cristatus* | 1791 | 766 |
| *Lissotriton helveticus* | 7047 | 2835 |

**Table 2. Description of the presence-only data used for each of nine species for calibration and cross-validation of habitat suitability models**. In the first part of the analyses, the model was calibrated with 70% of presence-only data and 30% of the data left were used for cross-validation.

#### *1.2. Standardised detection-nondetection data (external validation dataset)*

**Name of the citizen science program**: “Un Dragon dans mon Jardin”

**Coordination**: URCPIE – “Union régionale des centres d’initiatives pour l’environnement ».

For external SDM validation, we extracted detection-nondetection_amphibian data from a regional citizen science database. This database contained 576 monitored aquatic sites for the period 2013-2019, with observations made in the context of a programme aiming to estimate amphibian population trends (regionally called “Un Dragon dans mon Jardin”). Observers followed a standard protocol; each site was monitored three times separated by at least one month - one diurnal between January and March and two nocturnal between March and June – to cover different species’ breeding periods, during good weather conditions (no frost, no rain, no or weak wind). For each survey, three complementary methods were used to detect amphibians: an acoustic survey (5 min at 5 metres from the site without light) to detect breeding calls of male Anurans specie; an active visual survey using a flashlight torch (500-1000 lumens) to observe individuals and eggs and a catching survey using a net (3 net sweeps per site). These methods are commonly used for amphibian community surveys.

| | **CS.0** | | **VOL** | | **PRO** | |
|---|---|---|---|---|---|---|
| **Species** | Nb of DET | Nb of NoDET | Nb of DET | Nb of NoDET | Nb of DET | Nb of NoDET |
| **Anourans:** | | | | | | |
| *Bufo spinosus* | 79 | 195 | 31 | 93 | 25 | 87 |
| *Hyla arborea arborea* | 98 | 176 | 43 | 81 | 62 | 50 |
| *Pelodytes punctatus* | 19 | 255 | 7 | 117 | 20 | 92 |
| *Rana dalmatina* | 176 | 98 | 64 | 60 | 71 | 41 |
| *Rana temporaria* | 14 | 260 | 5 | 119 | 2 | 110 |
| **Urodeles:** | | | | | | |
| *Salamandra Salamandra terrestris* | 80 | 194 | 25 | 99 | 23 | 89 |
| *Triturus marmoratus* | 43 | 231 | 20 | 104 | 14 | 98 |
| *Triturus cristatus* | 30 | 244 | 16 | 108 | 24 | 88 |
| *Lissotriton helveticus* | 171 | 103 | 59 | 65 | 65 | 47 |

**Table 3. Description of the initial datasets without filtering for each of nine species used for external validation of habitat suitability models**. CS.0: all data from a citizen science program “Un Dragon dans mon jardin” without filter collected between 2013 and 2019; VOL: all additional data collected by volunteers in 2019; PRO: data collected by professionals. DET: 500m cells with detection of the species; NoDET: 500m nondetection-cells

| | CS.1 | | CS.2 | | CS.1 + ABS + SUP | | CS.2 + ABS + SUP | |
|---|---|---|---|---|---|---|---|---|
| | Nb of DET | Nb of NoDET | Nb of DET | Nb of NoDET | Nb of DET | Nb of NoDET | Nb of DET | Nb of NoDET |
| **Anourans:** | | | | | | | | |
| *Bufo spinosus* | 54 | 49 | 54 | 40 | 97 | 187 | 97 | 185 |
| *Hyla arborea* | 56 | 65 | 56 | 59 | 136 | 204 | 137 | 203 |
| *Pelodytes punctatus* | 16 | 63 | 15 | 65 | 40 | 249 | 40 | 211 |
| *Rana dalmatina* | 80 | 42 | 81 | 34 | 186 | 162 | 187 | 159 |
| *Rana temporaria* | 9 | 57 | 11 | 57 | 17 | 231 | 16 | 228 |
| **Urodeles:** | | | | | | | | |
| *Salamandra Salamandra* | 44 | 46 | 44 | 35 | 79 | 186 | 79 | 177 |
| *Triturus marmoratus* | 30 | 58 | 31 | 16 | 62 | 213 | 61 | 169 |
| *Triturus cristatus* | 17 | 71 | 19 | 21 | 52 | 241 | 53 | 198 |
| *Lissotriton helveticus* | 87 | 31 | 85 | 7 | 176 | 164 | 175 | 125 |

**Table 4. Description of the filtered datasets for each of nine species used for external validation of habitat suitability models**. CS.0: all data from a citizen science program "Un Dragon dans mon jardin" without filter collected between 2013 and 2019; SUP: all additional data collected by volunteers and by professionals in 2018-2019. CS.2 (or CS.1) + ABS (CS.2 (or CS.1) with 10% supplement absence cells in very unfavourable habitats). DET: 500m cells with detection of the species; NoDET: 500m nondetection-cells. Results for 1 interaction.

| | STRAT_CS | | STRAT_ALL | |
|---|---|---|---|---|
| | Nb data/strat for s2 | Nb data/strat for s3 | Nb data/strat for s2 | Nb data/strat for s3 |
| **Anourans:** | | | | |
| *Bufo spinosus* | 22 | 19 | 37 | 23 |
| *Hyla arborea arborea* | 17 | 11 | 42 | 23 |
| *Pelodytes punctatus* | 13 | 3 | 29 | 8 |
| *Rana dalmatina* | 18 | 19 | 38 | 30 |
| *Rana temporaria* | 9 | 10 | 8 | 15 |
| **Urodeles:** | | | | |
| *Salamandra Salamandra terrestris* | 19 | 21 | 23 | 25 |
| *Triturus marmoratus* | 9 | 10 | 13 | 16 |
| *Triturus cristatus* | 18 | 14 | 37 | 25 |
| *Lissotriton helveticus* | 18 | 13 | 26 | 13 |

**Table 5. Number of filtered data by stratification used for external validation of habitat suitability models (STRAT_CS and STRAT_ALL)**.

### *1.3. Sites selection for data complementation*

All supplementary sites (263 ponds without fish) were selected randomly in order to complete 2 landscape gradients: woody element (hedges + woods) density and pond density. The 132 ponds that we monitored were distributed in six 30x30 km sectors and gradients were complete in each sector. Sites were randomly sampled so as to decorrelate pond density and woody element density which are naturally dependant in our region. A seventh 30x30 km sector was sampled with volunteers during three sessions (see Figure 1 and 2). Other sampled ponds were selected throughout the region to complete the 2 landscapes gradient according to existing data from 2013 to 2018 (e.g. in Figure 3).

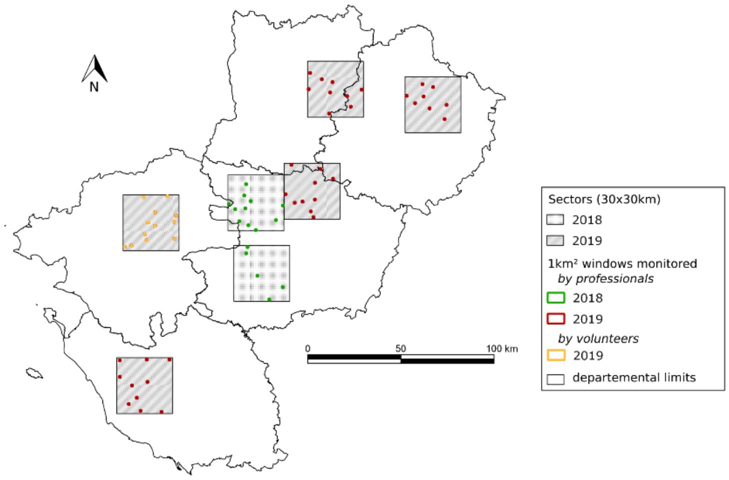


**Figure 1. Sectors of 30x30km monitored by professionals or volunteers in 2018 or 2019.** Gradients were complete in each sector and sites were randomly sampled so as to decorrelate pond density and woody element density. Three pounds without fish have been monitored in each windows of 1 km².

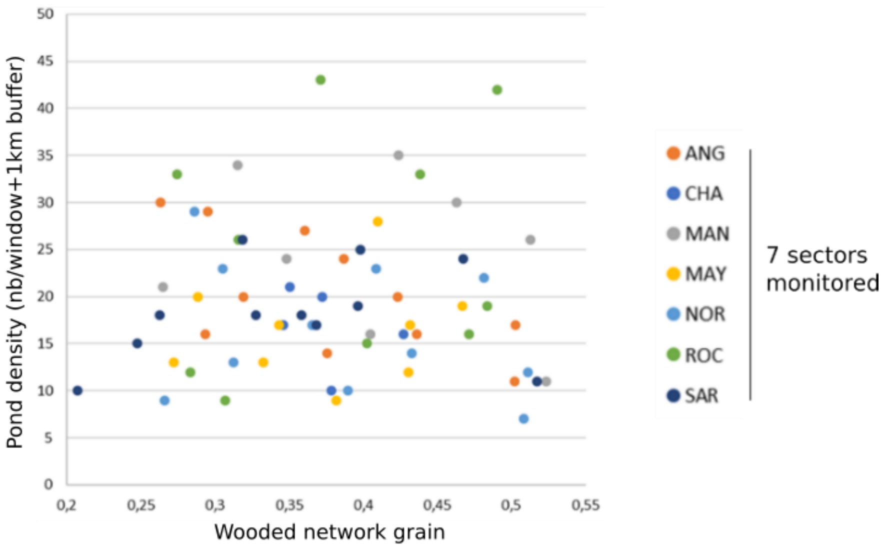


**Figure 2. Repartition of the monitored windows by professionals or volunteers in 2018 or 2019 in the 7 sectors (30x30km²) along pond density and wooded elements density.** Three pounds without fish have been monitored in each windows of 1 km². Volunteers monitored "SAR" sector and all others were monitored by professionals. Higher is "wooded network grain", lower is woody elements density.

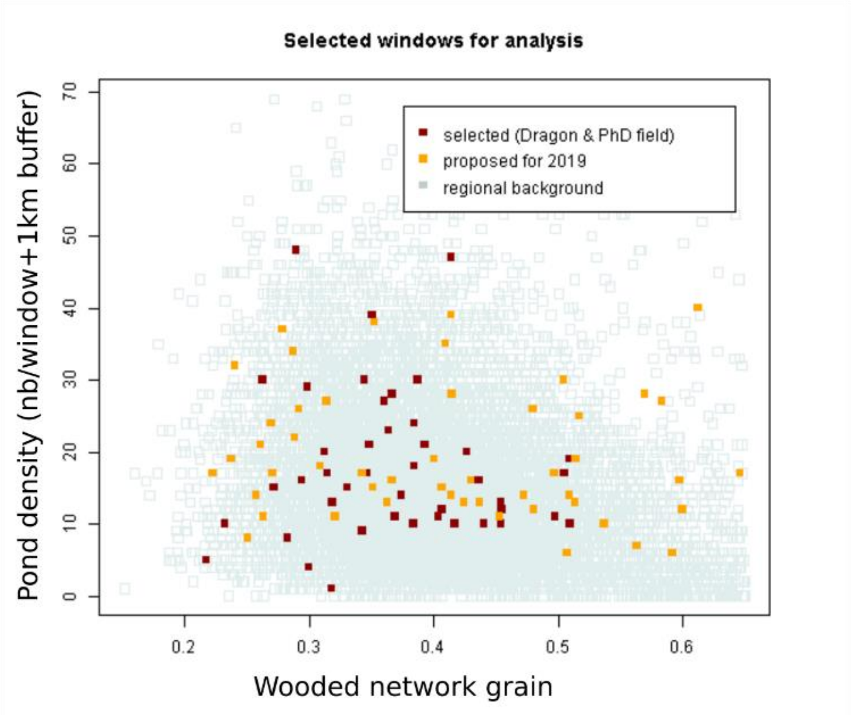


**Figure 3. Example of proposed windows for monitoring by volunteers and their distribution along the two gradients (pond density and woody element density).** Higher is “wooded network grain”, lower is woody elements density. “selected” data were existing data in 2018 after strong filtering and before additional field.

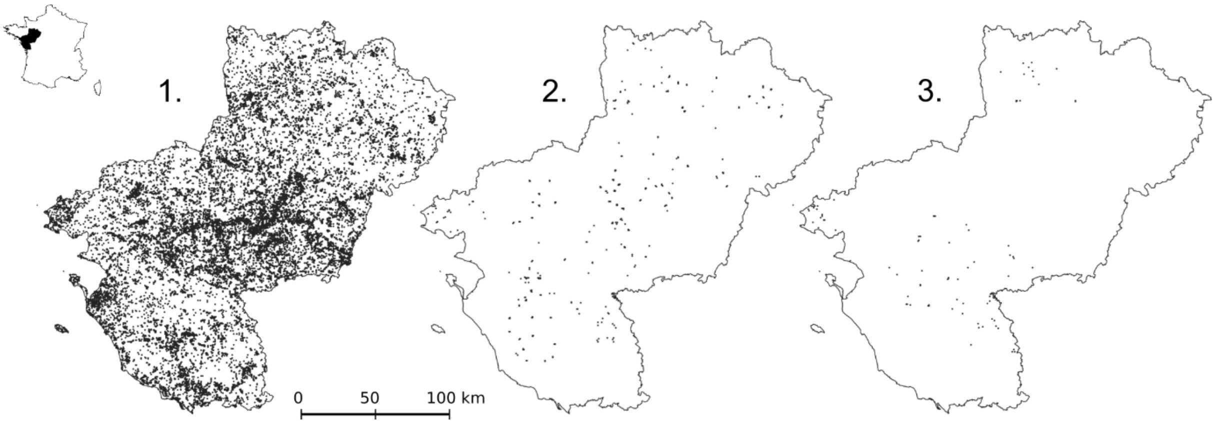


**Figure 4. Distribution of the 500m² cells with data for the opportunistic dataset and the external evaluation dataset (e.g. *CS.1+ABS+SUP*).** (1) All 500m² cells with at least one opportunistic observation (all species); (2) 500m² cells used as presence-absence data (with at least three surveys performed by an expert observer or six surveys by an intermediate observer) for external validation; (3) 500m² cells used only as presence if the species had been detected (sampling effort too weak for absence data) for external validation. The external dataset for validation is a compilation of (2) (presence-absence) and (3) (presence).

### 1.4. *Observation level and threshold values for the minimal sampling effort required to valid absence data for each species.*

Based on our observer classes, we set threshold values for the sampling effort needed to validate absence data (i.e. the minimal number of surveys called *N* and defined for each observer classes “expert”, ”intermediate” and “novice” called $N_{exp}$, $N_{int}$ and $N_{nov}$ respectively); absence data was validated when grid cells had been monitored by at least $N_{exp}$ nocturnal surveys conducted by an “expert” observer or at least $N_{int}$ nocturnal surveys by “intermediate” observer and at least $N_{nov}$ surveys by a "novice" observer. $N_{exp}$, $N_{int}$ and $N_{nov}$ were defined according to four species detection classes: species easily detected (e.g. *Rana dalmatina* and *Hyla arborea*) with $N_{exp}$=2 $N_{int}$=2 and $N_{nov}$=4 ; species with medium detection rate with $N_{exp}$=2, $N_{int}$=3 and no $N_{nov}$ (e.g. *Triturus cristatus* and *Lissotriton helveticus*) and species more difficult to detected with $N_{exp}$=3, $N_{int}$=4 and no $N_{nov}$ (e.g. *Triturus marmoratus*, *Salamandra salamandra*, *Bufo spinosus* and *Pelodytes punctatus*) or with $N_{exp}$=3, $N_{int}$=5 and no $N_{nov}$ (e.g. *Rana temporaria*). These classes were defined according to occupancy studies in France (Boissinot 2008 and Petitot et al., 2014) in Switzerland (Pellet et Schmidt 2005) and in UK with volunteers’ surveys (Sewell et al. 2010). Difference between observers’ groups were defined according to the species detection probability calculated for the monitoring methods used (i.e. acoustic, visual or direct sampling using a fishing net) by Boissinot 2008;

### 1.5. *Target species for absence validation (CS.2)*

| **Species** | Target species for absence validation |
|---|---|
| **Anourans:** | |
| *Bufo spinosus* | At least one other species detected |
| *Hyla arborea arborea* | At least one other species detected |
| *Pelodytes punctatus* | At least one other species detected |
| *Rana dalmatina* | At least one other species detected |
| *Rana temporaria* | At least one other species detected |
| **Urodeles:** | |
| *Salamandra Salamandra terrestris* | *Triturus cristatus or Triturus marmoratus or Lissotriton helveticus* |
| *Triturus marmoratus* | *Triturus cristatus* |
| *Triturus cristatus* | *Triturus marmoratus* |
| *Lissotriton helveticus* | At least one other species detected |

**Table 6. Target species used for absence validation for each studied species**
