## Appendix 2 for "How citizen science could improve Species Distribution Models and their independent assessment"

### Appendix 2 – Environmental variables: additional information

| Variable category | Code | Variable description | References |
|---|---|---|---|
| Climatic | CLIM_1 | first axis from a PCA on 12 worldclim variables and altitude (see Table 4) | Rothermel and Semlitsch, 2002; Pineda and Lobo, 2009; Girardello *et al.*, 2010; Hartel *et al.*, 2010 |
| | CLIM_2 | second axis from a PCA on 12 worldclim variables and altitude (see Table 4) | |
| Land cover | %WOOD_DM | Proportion of deciduous and mixed forest | Pope, Fahrig and Merriam, 2000; Pellet, Guisan and Perrin, 2004; Cushman, 2006; Zanini *et al.*, 2008; Hartel *et al.*, 2010; Boissinot *et al.*, 2015; Zhang *et al.*, 2016; Collins and Fahrig, 2017; Boissinot, Besnard and Lourdais, 2019 |
| | %WOOD_C | Proportion of coniferous forest | |
| | %CROP | Proportion of crop | Knutson *et al.*, 1999; Lehtinen, Galatowitsch and Tester, 1999; Mann *et al.*, 2009; Hartel *et al.*, 2010; Boissinot, Besnard and Lourdais, 2019 |
| | %PASTURE | Proportion of permanent pasture | Scribner *et al.*, 2001; Janin *et al.*, 2009; Hartel *et al.*, 2010 |
| | NB_PONDS | Ponds density | Janin *et al.*, 2009; Ribeiro *et al.*, 2011; Arntzen *et al.*, 2017; Boissinot, Besnard and Lourdais, 2019 |
| | L_HEDGE | Hedgerow density | Joly *et al.*, 2001; Pellet, Guisan and Perrin, 2004; Vos *et al.*, 2007; Angelone, Kienast and Holderegger, 2011; Boissinot, Besnard and Lourdais, 2019 |
| | L_ROAD_1ST | Primary Road density out of urban area | Vos and Chardon, 1998; Carr and Fahrig, 2001; Hels and Buchwald, 2001; Cushman, 2006; Eigenbrod, Hecnar and Fahrig, 2008; Hartel *et al.*, 2010; Petrovan and Schmidt, 2016; Boissinot, Besnard and Lourdais, 2019 |
| | L_ROAD_2ND | Secondary Road density out of urban area | |
| | L_RIVER | Permanent rivers and canals density | Ficetola *et al.*, 2008 |
| | %URBAN | Proportion of urban (build up) | Rubbo and Kiesecker, 2005; Gagné and Fahrig, 2007; Hartel *et al.*, 2010; Cayuela *et al.*, 2015; Zhang *et al.*, 2016 |

**Table 1. Environmental variables used for species distribution modelling of each amphibian species in Pays de la Loire region and associated references.** See Figure 2/ table 4 for the list of climatic variables.

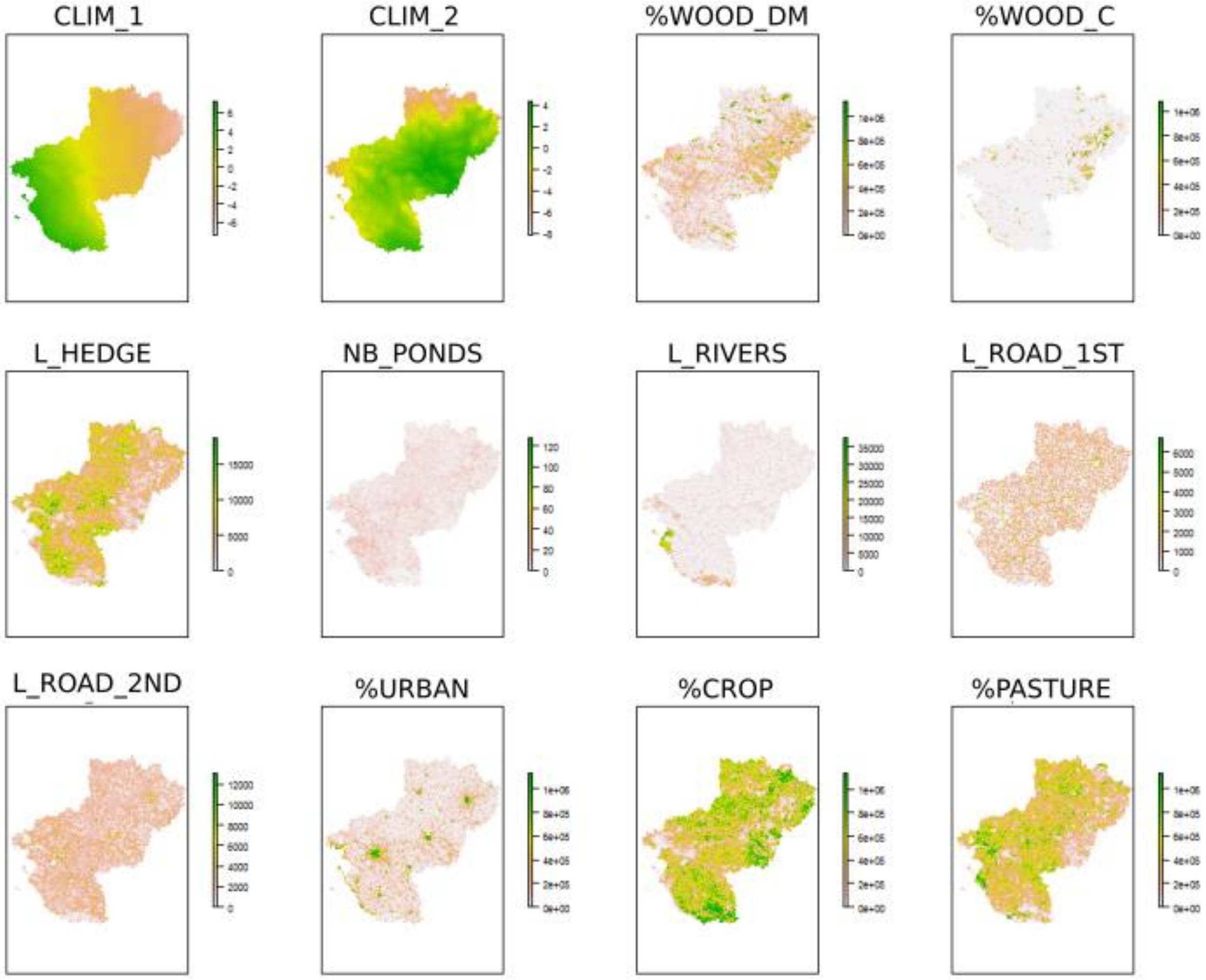


**Figure 1. Environmental variables used for modelling species distribution** (before scaling). Final maps resolution: 500m. CLIM_1 and CLIM_2 initial resolution was 5km².

| Variables | VIF |
|---|---|
| CLIM_1 | 1,26 |
| CLIM_2 | 1,09 |
| %WOOD_DM | 2,86 |
| %WOO8C | 1,89 |
| L_HEDGE | 1,89 |
| NB_PONDS | 1,18 |
| L_RIVERS | 1,27 |
| L_ROAD_1ST | 1,05 |
| L_ROAD_2ND | 1,18 |
| %URBAN | 2,66 |
| %CROP | 5,26 |
| %PASTURE | 4,67 |

**Table 2. Variance inflation factor values for each predictor**

| | CLIM_1 | CLIM_2 | %WOOD_DM | %WOO8C | L_HEDGE | NB_PONDS | L_RIVERS | L_ROAD_1ST | L_ROAD_2ND | %URBAN | %CROP | %PASTURE |
|---|---|---|---|---|---|---|---|---|---|---|---|---|
| **CLIM_1** | 1,00 | | | | | | | | | | | |
| **CLIM_2** | 0,00 | 1,00 | | | | | | | | | | |
| **%WOOD_DM** | -0,19 | 0,08 | 1,00 | | | | | | | | | |
| **%WOO8C** | -0,13 | 0,07 | 0,31 | 1,00 | | | | | | | | |
| **L_HEDGE** | 0,14 | -0,19 | -0,33 | -0,28 | 1,00 | | | | | | | |
| **NB_PONDS** | 0,15 | 0,03 | -0,10 | -0,09 | 0,27 | 1,00 | | | | | | |
| **L_RIVERS** | 0,28 | -0,03 | -0,10 | -0,06 | -0,09 | 0,03 | 1,00 | | | | | |
| **L_ROAD_1ST** | -0,06 | 0,01 | -0,06 | -0,05 | 0,04 | 0,06 | -0,06 | 1,00 | | | | |
| **L_ROAD_2ND** | 0,08 | 0,04 | -0,20 | -0,19 | 0,10 | 0,09 | -0,11 | 0,08 | 1,00 | | | |
| **%URBAN** | 0,15 | 0,08 | -0,06 | -0,08 | -0,16 | 0,02 | -0,03 | 0,12 | 0,25 | 1,00 | | |
| **%CROP** | -0,08 | 0,07 | -0,44 | -0,30 | 0,01 | -0,14 | -0,14 | 0,04 | 0,08 | -0,31 | 1,00 | |
| **%PASTURE** | 0,09 | -0,16 | -0,28 | -0,25 | 0,57 | 0,27 | 0,18 | -0,02 | 0,04 | -0,22 | -0,38 | 1,00 |

**Table 3. Pearson correlation test for each predictor**

### Climatic variables

| | | | Contribution (ACP) | | | | |
|---|---|---|---|---|---|---|---|
| | | | Dim.1 | Dim.2 | Dim.3 | Dim.4 | Dim.5 |
| **VARIABLES** | Altitude | r_alt | 7,18 | 7,52 | 0,62 | 44,81 | 22,89 |
| | Annual mean temperature (°C) | r_bio1 | 10,91 | 5,80 | 4,91 | 1,11 | 2,24 |
| | Mean temperature of warmest quarter (°C) | r_bio10 | 2,47 | 16,01 | 18,18 | 12,75 | 1,48 |
| | Mean temperature of coldest quarter (°C) | r_bio11 | 14,67 | 0,66 | 0,02 | 1,48 | 1,88 |
| | Annual precipitation (mm) | r_bio12 | 9,50 | 7,44 | 0,14 | 6,53 | 20,79 |
| | Precipitation seasonality (C of V) | r_bio15 | 14,29 | 0,52 | 1,82 | 5,94 | 0,83 |
| | Precipitation of wettest quarter (mm) | r_bio16 | 11,82 | 4,32 | 0,01 | 6,77 | 11,86 |
| | Precipitation of driest quarter (mm) | r_bio17 | 1,10 | 18,31 | 14,72 | 2,48 | 0,17 |
| | Precipitation of warmest quarter (mm) | r_bio18 | 0,06 | 16,39 | 43,64 | 13,22 | 0,66 |
| | Max temperature of warmest week (°C) | r_bio5 | 1,95 | 18,28 | 11,16 | 2,30 | 6,93 |
| | Temperature annual range (Bio05-Bio06) (°C) | r_bio7 | 12,01 | 4,07 | 4,43 | 1,96 | 5,07 |
| | Mean temperature of wettest quarter (°C) | r_bio8 | 14,03 | 0,67 | 0,34 | 0,64 | 25,19 |

**Table 4. variable selection and their contribution for each axis of the PCA**

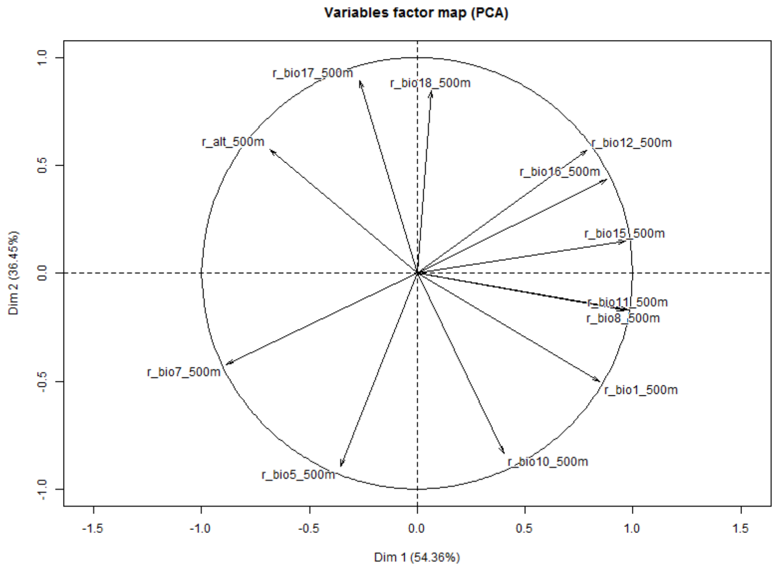


**Figure 2. Representation of the attributes on the factors 1 and 2 obtain by PCA by descriptor**. We use layer for Worldclim at regional extent to obtain 2 layers according to axis 1 (CLIM_1) and axis 2 (CLIM_2) of the ACP. Final resolution was 500m to homogenize pixels size of climatic variables and landscape variables layers.
