## Appendix 3 for "How citizen science could improve Species Distribution Models and their independent assessment"

**Appendix 3 – Definition and modelling of sampling effort for SDM at regional extent**

Presence-only data are abundant but have poor quality, doesn't give any information about absence-data, have few metadata and come from different sources (Robinson et al., 2020). To get true absence data need a higher sampling effort than presence data (Graham et al., 2004). In order to overcome the problem of missing absence data needed for most SDM, pseudo-absence selection strategies have been developed to select absence data where real absence is most likely (Phillips et al., 2009; Barbet-Massin et al., 2012). Understanding the structure and intensity of sampling effort in space is essential to determine whether an undetected species is truly absent. For example, it may be conditioned by site accessibility (Kadmon, Farber & Danin, 2004; Phillips et al., 2009), site attractiveness or observer distribution (Phillips et al., 2009; Robinson, Ruiz-Gutierrez & Fink, 2018).

Here we considered three main sources of bias in pseudo-absence selection: **accessibility**, linked to distance from roads or urban areas (Kadmon, Farber & Danin, 2004; Barbet-Massin et al., 2012b), **attractiveness**, relating to oversampling in protected sites or natural tourist sites (Robinson, Ruiz-Gutierrez & Fink, 2018) and **observer distribution and activity**, because certain administrative areas are covered by particularly active nature protection organisations.

### 1. Definitions

#### 1.1. Accessibility (ACCESS)

*Hypothesis:*

- More accessible is the site, higher is the probability that it has been sampled
- Accessibility is conditioned by the presence of a road or path.

Secondary roads and paths are probably a better indicator of accessibility than primary roads. Indeed, it is impossible to stop at a highway or a dual carriageway roadside to access a site. It is also generally difficult to stop at of a primary roadside (parking prohibition and / or heavy traffic and / or difficult parking). In addition, direct observations on these road types is very difficult and dangerous. Direct observation of individuals on the roads (traveling individual or carcass) is easier on secondary roads because of lower nighttime traffic and lower speed.

Hence, **we defined accessibility as the distance to secondary roads and paths as well as indirectly as the distance from an urban area** (there is a high road density in urban area)**,** as an element facilitating access.

*Modelling*

We used a half-normal function to describe sampling probability according to distance from secondary road and paths. The probability density function of the half-normal is given by:

$$f(x;a) = \frac{\sqrt{2}}{a\sqrt{\pi}}\exp(-\frac{x^2}{2a^2}) \qquad (1)$$

Where a is a constant corresponding to the x value of the inflection point. We use a=100m which means that the site is twice as likely to be sampled close to the road than at a distance of 100m and thereafter probability strongly declines with distance. This is the probability function commonly used in distance-sampling methods (Buckland et al. 1993).

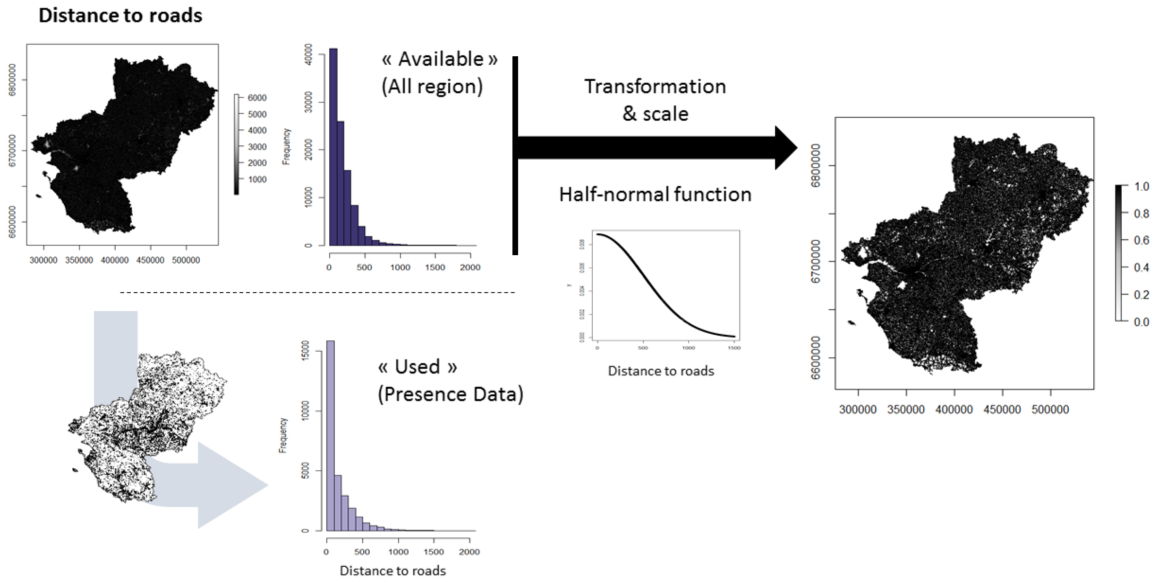


**Figure 1. Transformation and scale of "distance to road" layer with a half-normal function (100m resolution).**

#### 1.2. <u>Observer distribution and activity (OBS)</u>

*<u>Hypothesis:</u>*

- Sampling effort is higher where observer density is higher, e.g. close to main cities and university structures (OBS1);
- Observer activity and data collection access is higher where nature protection organisations are more active (OBS2).

These sources of bias are especially important in the case of "Maine-et-Loire" administrative area, which contains 38% of the dataset. To consider this imbalance between administrative areas, we sampled pseudo-absences in each according to the proportion of existing data (OBS2). Distances to a major city were converted to probability distributions using a half-normal function with a=50000m (OBS1).

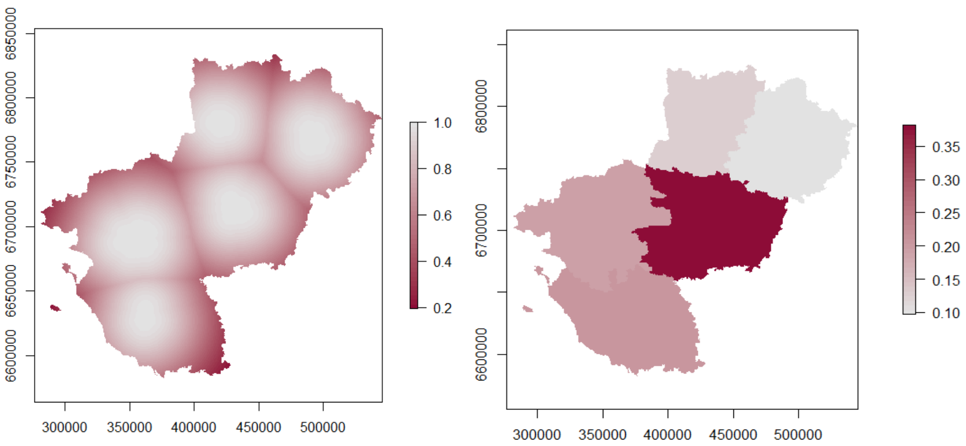

**Figure 2. Observer distribution and activity layers (left: OBS1; right: OBS2)**

#### 1.3. Site attractiveness (ATTRACT)

*Hypothesis:*

- Sampling effort is higher where sites are attractive for naturalists
- Naturalist motivation to go on the field and collect data might be led to the natural beauty of the area, the localisation of naturalist hotspot or protected area or political interest for nature protection.

In Pays-de-la-Loire region, most protected areas and natural tourist sites are located near the Loire and Maine rivers. Therefore, we used **the distance to the Loire and Maine rivers as a proxy for site attractiveness**.

*Modelling*

We used a half-normal function for describing sampling probability according to distance from Loire river with a=4000m (inflection point). 4000m corresponds to twice the maximum distance of any protected area from the Loire river.

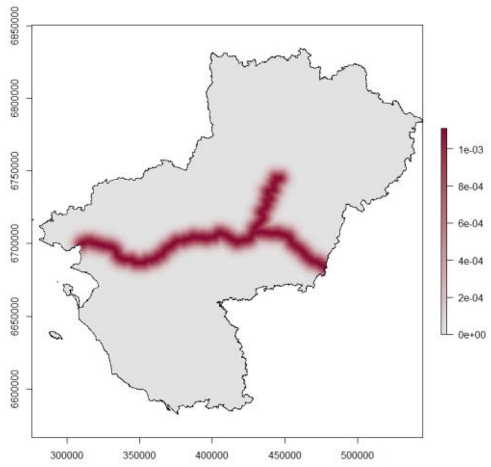


**Figure 3. Site attractiveness near two main rivers: half-normal transformation (ATTRACT)**

### 2. Compilation

We considered the interaction between attractiveness (distance to the 2 main rivers) and distance to cities may explain additional sampling effort heterogeneity. We obtained a final layer using the following relationship:

**(ATTRACT + OBS1)*ACCESS*OBS2**

With:

**ATTRACT**: Distance to the two main rivers (indirectly related to main protected areas and touristic natural places)

**OBS1**: Distance to the main cities

**ACCESS**: Distance to the secondary roads and paths

**OBS2**: local nature protection organisations activity (indirectly related to the proportion of observations by administrative area)

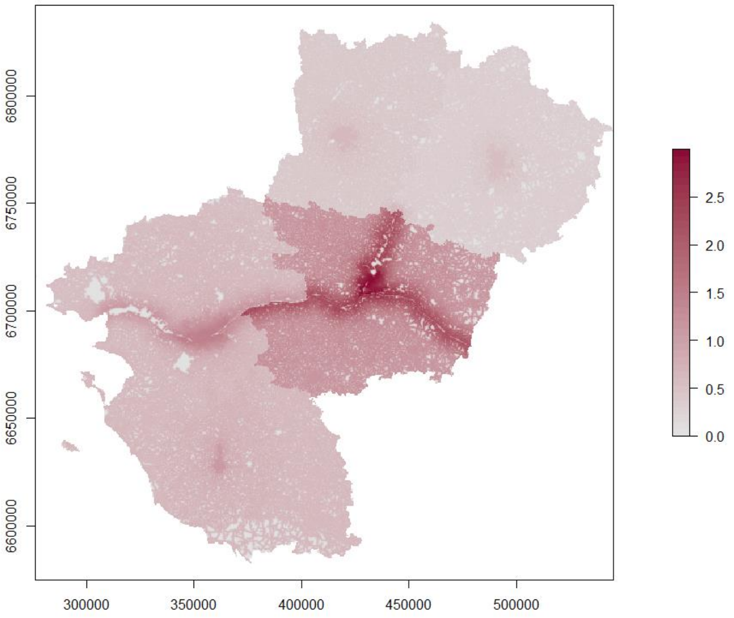


**Figure 4. Final layer used to include sampling effort heterogeneity in pseudo-absence sampling.**

**<u>Bibliography</u>**

Barbet-Massin, M., Jiguet, F., Albert, C. H. & Thuiller, W. (2012a) 'Selecting pseudo-absences for species distribution models: how, where and how many?', Methods in Ecology and Evolution, 3(2), pp. 327–338. doi: 10.1111/j.2041-210X.2011.00172.x.

Barbet-Massin, M., Jiguet, F., Albert, C. H. & Thuiller, W. (2012b) 'Selecting pseudo-absences for species distribution models: How, where and how many?', Methods in Ecology and Evolution, 3(2), pp. 327–338. doi: 10.1111/j.2041-210X.2011.00172.x.

Graham, C. H., Ferrier, S., Huettman, F., Moritz, C. & Peterson, A. T. (2004) 'New developments in museum-based informatics and applications in biodiversity analysis', Trends in Ecology and Evolution, 19(9), pp. 497–503. doi: 10.1016/j.tree.2004.07.006.

Kadmon, R., Farber, O. & Danin, A. (2004) 'Effect of roadside bias on the accuracy of predictive maps produced by bioclimatic models', Ecological Applications, 14(2), pp. 401–413. doi: 10.1890/02-5364.

Phillips, S. J. … Ferrier, S. (2009) 'Sample selection bias and presence-only distribution models: Implications for background and pseudo-absence data', Ecological Applications, 19(1), pp. 181–197. doi: 10.1890/07-2153.1.

Robinson, O. J., Ruiz-Gutierrez, V., Reynolds, M. D., Golet, G. H., Strimas-Mackey, M. & Fink, D. (2020) 'Integrating citizen science data with expert surveys increases accuracy and spatial extent of species distribution models', Diversity and Distributions, (October 2019), p. 806547. doi: 10.1101/806547.

Robinson, O. J., Ruiz-Gutierrez, V. & Fink, D. (2018) 'Correcting for bias in distribution modelling for rare species using citizen science data', Diversity and Distributions, 24(4), pp. 460–472. doi: 10.1111/ddi.12698.
