## Appendix 4 for "How citizen science could improve Species Distribution Models and their independent assessment"

### Appendix 4 – supplementary results

**1.**

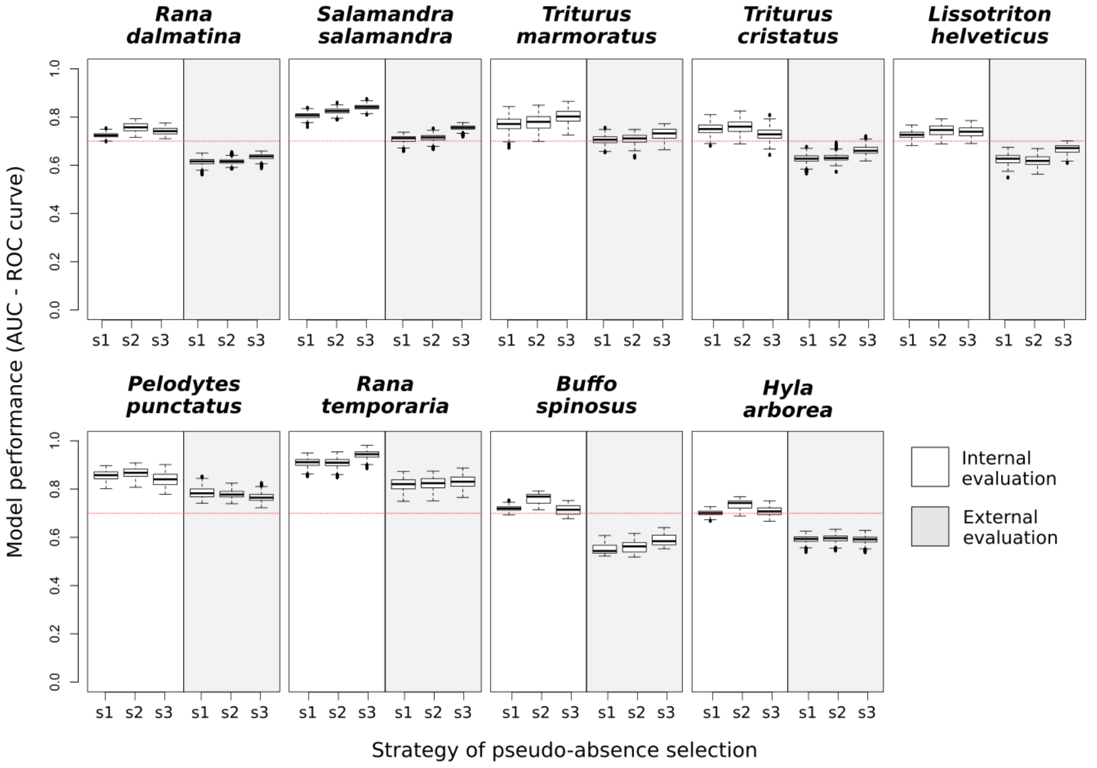


**2.**

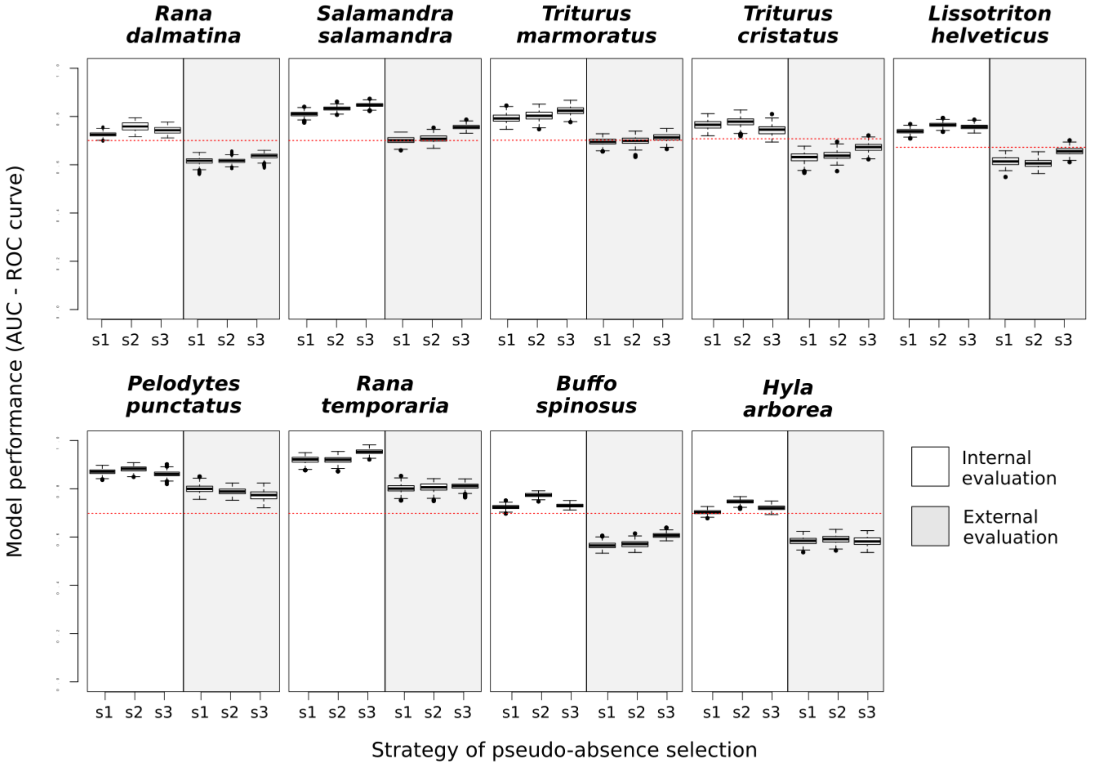


**Figure 1. Models performance for the 9 studied species assessed by external or internal evaluation according to different pseudo-absence selection strategies including or not sampling effort**. Evaluation by AUC under the ROC for GAM (1) and Random Forest (2). Artificial absence sampling strategies are s1 (random background points selection), s2 (random pseudo-absence selection excluding known presence points) and s3 (random pseudo-absence selection excluding known

presence points and adjusted to consider sampling effort). By strategy, 10 replicates of the artificial absence points generation processes with 30 iterations each with 70% of the data. Black dotted line indicates the 0.70 threshold above witch models have an acceptable level of accuracy. Boxplots showing upper whisker (maximum data point), interquartile range box (top line = 75% of the data ≤ this value; middle line = median; lower line = 25% of the data ≤ this value) and lower whisker (minimum data point). Outliers are shown by dots.

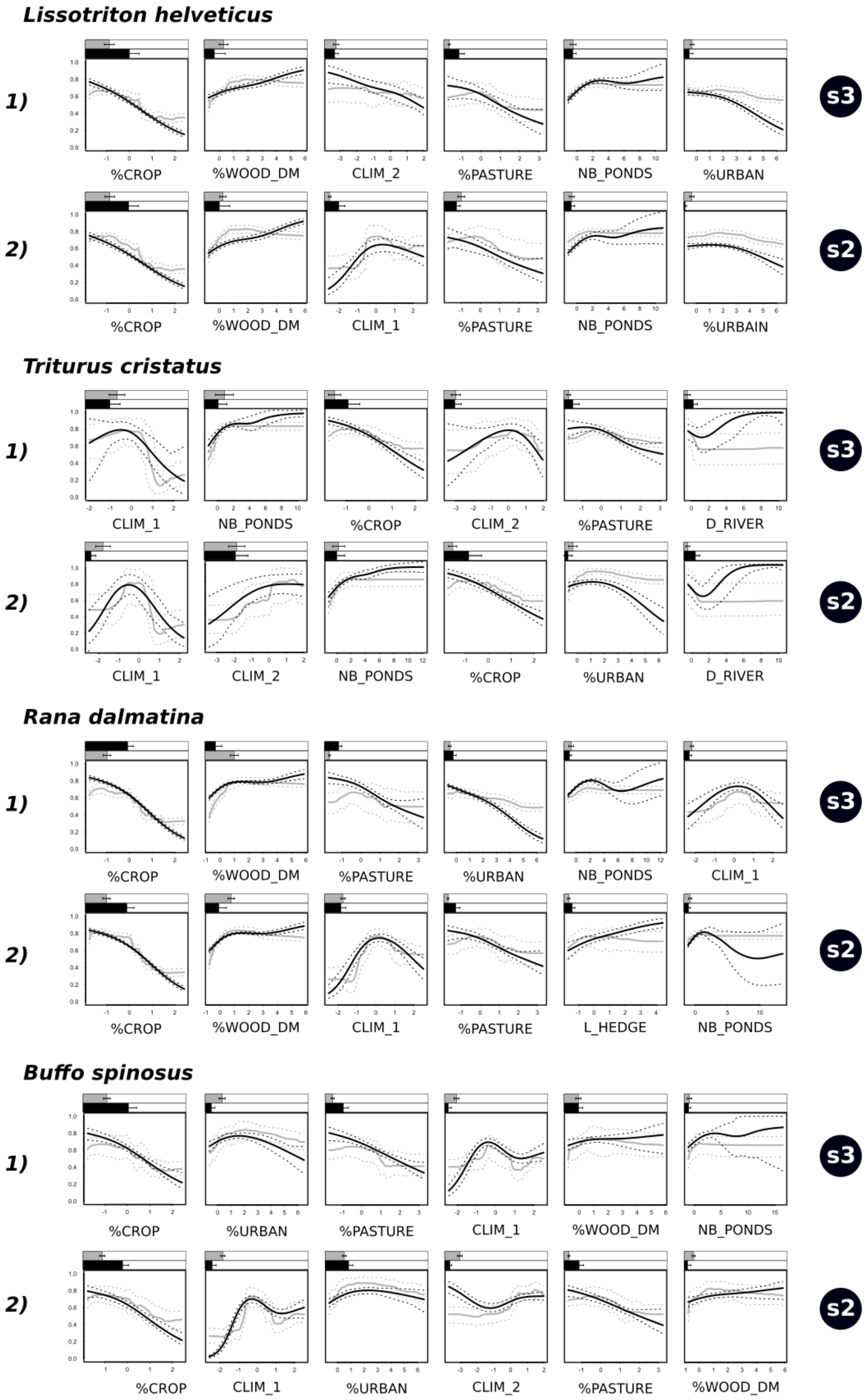


**Figure 2. Response curves for models selected by internal or external evaluation for four generalist species showing difference in model selection according to evaluation dataset used.** Results shown are for GAM (black line) and RF (grey line). Associate dotted lines show maximal range of all iterations

(50 iteration each 10 replicates). Bars above responses curves show mean percentage importance of variables and standard error (GAM in black and RF in grey). 1) best model with external evaluation; 2) best model with internal evaluation. s2 (random pseudo-absence selection excluding known presence points) and s3 (random pseudo-absence selection adjusted to consider site accessibility and sampling effort). For description of variables, see Table 2.

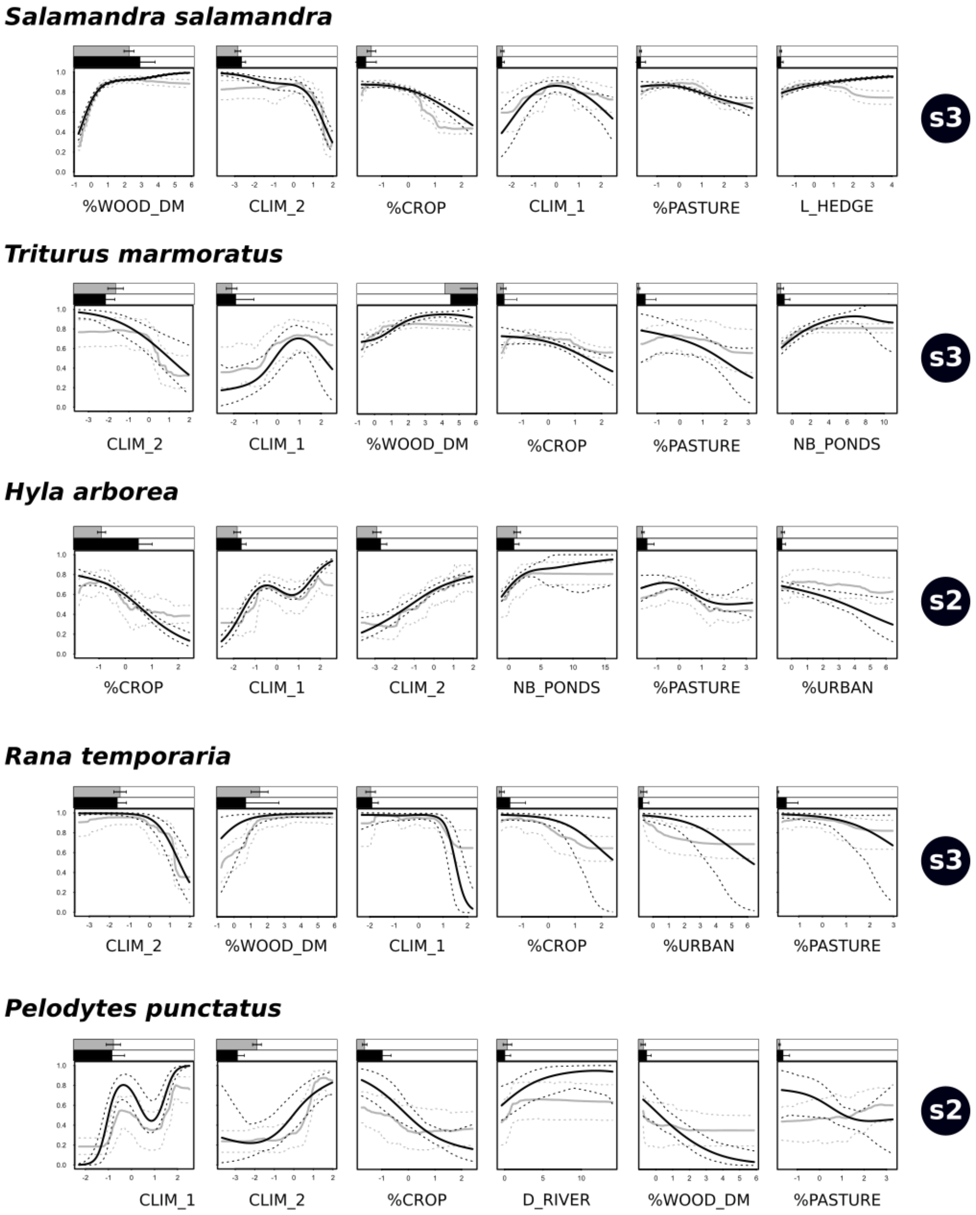


**Figure 3. Response curves for models selected by internal or external evaluation for five species showing no difference in model selection according to evaluation dataset used.** Best model with external evaluation and internal evaluation. Results shown are for GAM (black line) and RF (grey line). Associate dotted lines show maximal range of all iterations (50 iteration each 10 replicates). Bars above responses curves show mean percentage importance of variables and standard error (GAM in black and RF in grey). s2 (random pseudo-absence selection excluding known presence points) and s3 (random pseudo-absence selection adjusted to consider site accessibility and sampling effort). For description of variables, see Table 2.

| | | *Salamandra salamandra* | | | *Triturus marmoratus* | | | *Triturus cristatus* | | | *Lissotriton helveticus* | | | *Rana dalmatina* | | | *Hyla arborea* | | | *Rana temporaria* | | | *Pelodytes punctatus* | | | *Bufo spinosus* | | |
|---|---|---|---|---|---|---|---|---|---|---|---|---|---|---|---|---|---|---|---|---|---|---|---|---|---|---|---|---|
| | | **SEN** | **SPE** | **AUC** | **SEN** | **SPE** | **AUC** | **SEN** | **SPE** | **AUC** | **SEN** | **SPE** | **AUC** | **SEN** | **SPE** | **AUC** | **SEN** | **SPE** | **AUC** | **SEN** | **SPE** | **AUC** | **SEN** | **SPE** | **AUC** | **SEN** | **SPE** | **AUC** |
| **CS.0** | *s2* | 0,64 | 0,56 | 0,65 | 0,63 | 0,41 | 0,58 | **0,90** | 0,56 | 0,79 | **0,69** | 0,29 | 0,53 | **0,68** | 0,39 | **0,57** | **0,82** | 0,43 | 0,68 | 0,71 | **0,87** | 0,87 | 0,53 | 0,83 | 0,73 | **0,62** | **0,33** | 0,52 |
| | *s3* | **0,73** | 0,56 | **0,70** | **0,65** | **0,45** | **0,61** | 0,87 | **0,61** | 0,79 | 0,64 | **0,34** | 0,54 | 0,60 | **0,45** | 0,54 | 0,78 | **0,49** | 0,68 | **0,86** | 0,84 | 0,88 | **0,68** | 0,84 | 0,74 | 0,59 | 0,30 | **0,54** |
| **CS.1** | s2 | 0,65 | 0,61 | 0,67 | 0,60 | 0,57 | 0,62 | **0,87** | 0,49 | 0,75 | **0,67** | 0,26 | 0,52 | **0,57** | 0,38 | 0,52 | **0,71** | 0,48 | 0,64 | 0,65 | **0,86** | 0,83 | 0,49 | 0,81 | 0,68 | 0,59 | **0,34** | 0,52 |
| | s3 | **0,73** | **0,63** | **0,74** | **0,58** | **0,60** | **0,65** | 0,84 | **0,53** | 0,75 | 0,62 | **0,37** | **0,55** | 0,53 | **0,44** | 0,52 | 0,68 | **0,52** | 0,64 | **0,82** | 0,81 | 0,84 | **0,65** | 0,82 | **0,70** | 0,60 | 0,32 | **0,54** |
| **CS.2** | s2 | 0,65 | 0,59 | 0,67 | 0,59 | 0,67 | 0,72 | **0,87** | 0,53 | **0,78** | **0,66** | 0,33 | 0,61 | **0,57** | 0,40 | 0,53 | **0,71** | 0,50 | 0,66 | 0,64 | **0,85** | 0,83 | 0,49 | 0,82 | 0,68 | 0,60 | 0,32 | 0,52 |
| | s3 | **0,73** | **0,62** | **0,75** | 0,58 | **0,82** | 0,72 | 0,85 | 0,55 | 0,75 | 0,62 | **0,60** | **0,63** | 0,53 | **0,46** | 0,52 | 0,68 | **0,53** | 0,66 | **0,82** | 0,81 | 0,84 | **0,65** | 0,82 | **0,71** | 0,61 | 0,31 | **0,54** |
| **CS.1 + ABS** | s2 | 0,66 | 0,70 | 0,74 | 0,59 | 0,66 | 0,69 | **0,87** | **0,58** | 0,79 | **0,66** | 0,52 | 0,65 | **0,57** | 0,58 | 0,65 | **0,71** | 0,59 | 0,71 | 0,64 | **0,88** | 0,86 | 0,49 | 0,73 | 0,65 | 0,60 | 0,40 | 0,54 |
| | s3 | **0,73** | **0,72** | **0,80** | 0,58 | **0,69** | **0,71** | 0,84 | 0,62 | 0,79 | 0,62 | **0,61** | **0,71** | 0,53 | **0,62** | 0,65 | 0,68 | **0,63** | 0,72 | **0,82** | 0,84 | 0,87 | **0,65** | **0,78** | **0,70** | 0,60 | **0,45** | **0,56** |
| **CS.2 + ABS** | *s2* | 0,66 | 0,69 | 0,74 | 0,59 | 0,78 | 0,81 | **0,88** | **0,64** | **0,83** | **0,67** | 0,76 | 0,80 | **0,57** | 0,61 | 0,67 | **0,71** | 0,61 | 0,73 | 0,64 | **0,88** | 0,86 | 0,48 | 0,76 | 0,66 | 0,59 | 0,39 | 0,54 |
| | *s3* | **0,73** | **0,72** | **0,80** | 0,58 | **0,87** | 0,81 | 0,84 | 0,65 | 0,81 | 0,62 | **0,88** | **0,86** | 0,53 | **0,64** | 0,67 | 0,68 | **0,64** | 0,74 | **0,82** | 0,84 | 0,86 | **0,64** | **0,79** | **0,70** | 0,60 | **0,45** | **0,56** |
| **CS.2 + ABS + SUP** | *s2* | 0,68 | 0,84 | 0,75 | 0,58 | **0,82** | 0,82 | 0,70 | 0,52 | 0,65 | **0,51** | 0,77 | 0,68 | **0,53** | 0,66 | 0,65 | **0,63** | **0,56** | 0,63 | 0,59 | **0,91** | 0,85 | **0,69** | 0,75 | 0,80 | 0,48 | 0,59 | 0,57 |
| | *s3* | **0,73** | 0,84 | **0,80** | **0,62** | 0,78 | 0,81 | **0,73** | 0,53 | **0,68** | 0,49 | 0,78 | **0,74** | 0,51 | **0,70** | **0,68** | 0,57 | 0,54 | 0,63 | **0,76** | 0,86 | 0,85 | 0,63 | **0,81** | 0,80 | 0,48 | **0,69** | **0,63** |
| **PRO** | *s2* | 0,42 | 0,88 | 0,76 | 0,58 | **0,83** | **0,85** | 0,66 | 0,43 | 0,56 | **0,41** | 0,76 | 0,61 | **0,51** | 0,62 | 0,58 | **0,59** | **0,39** | 0,51 | 0,50 | 0,95 | 0,71 | **0,81** | 0,74 | 0,85 | **0,37** | 0,74 | 0,57 |
| | *s3* | **0,45** | 0,87 | **0,78** | **0,79** | 0,77 | 0,83 | **0,71** | **0,46** | **0,61** | 0,37 | **0,79** | **0,64** | 0,48 | **0,75** | **0,63** | 0,51 | 0,33 | **0,54** | 0,50 | 0,92 | 0,71 | 0,62 | **0,81** | 0,84 | 0,29 | **0,82** | **0,63** |
| **STRAT_CS** | s2 | 0,69 | 0,67 | 0,74 | 0,71 | **0,72** | **0,81** | **0,73** | 0,58 | 0,71 | 0,67 | 0,81 | 0,82 | **0,64** | **0,65** | **0,73** | 0,72 | 0,64 | 0,73 | 0,79 | 0,57 | 0,69 | 0,80 | 0,58 | 0,81 | 0,56 | 0,54 | 0,60 |
| | s3 | **0,72** | 0,68 | **0,79** | 0,70 | 0,69 | 0,78 | 0,70 | 0,57 | 0,70 | 0,68 | 0,82 | **0,88** | 0,62 | 0,62 | 0,70 | 0,71 | 0,63 | **0,75** | **0,95** | **0,60** | **0,79** | **0,92** | **0,75** | **0,93** | **0,63** | **0,58** | **0,66** |
| **STRAT_ALL** | s2 | 0,74 | 0,67 | 0,76 | **0,80** | **0,74** | **0,87** | 0,74 | 0,56 | 0,70 | 0,66 | 0,76 | 0,78 | 0,62 | **0,65** | **0,70** | **0,68** | **0,62** | 0,70 | 0,84 | 0,57 | 0,73 | 0,78 | 0,58 | 0,76 | 0,58 | 0,55 | 0,62 |
| | s3 | 0,73 | 0,66 | 0,77 | 0,78 | 0,70 | 0,83 | **0,78** | 0,57 | **0,73** | **0,71** | **0,83** | **0,89** | 0,62 | 0,63 | 0,69 | 0,66 | 0,60 | **0,72** | **0,96** | 0,58 | **0,81** | **0,83** | **0,61** | **0,80** | **0,67** | **0,59** | **0,68** |

**Table 2. Model performance according to different filters and complementation fieldwork on the external evaluation set.** External sets used are: *CS.0* (all data from the standardized citizen science dataset – i.e. from 576 sites projected on 276 grind cells); *CS.1* (with a sampling effort threshold for absence selection and minimum of 1km distance between 500m grind cells); *CS.2* (as CS.1 adding the use of target species for absence selection); *CS.2 (or CS.1) + ABS* (*CS.2 (or CS.1)* with 10% supplement absence cells in very unfavourable habitats); *PRO* (data collected by professionals only in 2018-2019); *CS.2 + ABS + SUP* (citizen science data cited before adding all complementary fieldwork by professionals and volunteers); STRAT_CS (stratified data selection from CS.2+ABS with complementary field by volunteers); STRAT_ALL (stratified data selection from CS.2+ABS+SUP). Models assessed: s2 (random pseudo-absence selection excluding known presence points) and s3 (random pseudo-absence selection constrained to account sampling effort and correct sampling bias). SEN: sensitivity, SPE: specificity. Bold values show best values between s2 et s3 with delta > 0.02 and grey cells show species with less than 2 presence data. All analyses with a random sampling in presence selection with a distance condition or a stratified random selection were performed using 100 bootstraps (mean calculation).
